# Crosstalk of noradrenergic Ca^2+^ and cAMP signaling in astrocytes of the murine olfactory bulb

**DOI:** 10.1101/2025.11.10.687535

**Authors:** Jessica Sauer, Antonia Beiersdorfer, Franz Lennart Schmidt, Mathias Nordbeck, Oana Constantin, Daniela Hirnet, Christine E. Gee, Christian Lohr

**Affiliations:** Division of Neurophysiology, University of Hamburg, Germany; Center for Translational Neuromedicine, University of Copenhagen, Denmark; Sorbonne Université, Institut du Cerveau - Paris Brain Institute - ICM, Inserm, CNRS, APHP, Hôpital de la Pitié Salpêtrière, Paris, France; Institute of Synaptic Physiology, ZMNH, University Medical Center Hamburg-Eppendorf, Germany

**Keywords:** Norepinephrine, adenylyl cyclase, phenylephrine, cyclic adenosine monophosphate, Ca^2+^, astrocytes

## Abstract

Cyclic adenosine monophosphate (cAMP) and Ca^2+^ are ubiquitous second messengers that regulate gene expression, metabolism, and synaptic plasticity. Here, we identified a complex interplay between Ca^2+^ and cAMP signaling pathways in mouse olfactory bulb astrocytes. Norepinephrine (NE) elevated both Ca^2+^ and cAMP levels via α_1_ and α_2_ adrenergic receptors, whereas β receptors triggered only cAMP responses. The α_1_ receptor agonist phenylephrine increased cAMP, but this effect was suppressed when Ca^2+^ elevations were blocked by Ca^2+^ depletion and removal of external Ca_2+_. We found that α_1A_ and α_1D_ receptors are key targets for phenylephrine, acting through Ca_2+_/calmodulin-dependent adenylyl cyclases AC1 and AC3 downstream of α_1_ receptor activation. Moreover, α_2_ receptor stimulation raised Ca^2+^ levels, thereby stimulating cAMP production, yet also reduced forskolin-induced cAMP elevations, indicating that α_2_ receptors can both inhibit adenylyl cyclase via G_i_ and stimulate AC1/AC3 via Ca^2+^ signaling. Together, these findings reveal intricate crosstalk between noradrenergic Ca^2+^ and cAMP signaling in olfactory bulb astrocytes mediated by all three adrenergic receptor subtypes.

## Introduction

The locus coeruleus (LC) serves as the primary source of norepinephrine (NE) in the brain, where NE is synthesized in noradrenergic neurons by beta-hydroxylation of dopamine (Fuller, 1982). Notably, approximately 40 % of all LC neurons project to the olfactory bulb (OB), a number that is tenfold greater than projections to any other region of the cerebral cortex (Shipley et al., 1985). NE and its effects on neural dynamics in the OB are essential for the formation of stable and specific olfactory memories (Linster et al., 2020). NE enhances the contrast of weak neuronal sensory inputs in the OB by modulating mitral cell responses (Linster et al., 2011). The NE system plays a pivotal role in both health and disease. In Parkinsońs Disease, for instance, not only does the substantia nigra degenerate, but so does the LC, with olfactory dysfunction being one of the early symptoms (Paredes-Rodriguez et al., 2020). In addition, dysfunction of astrocytes is also implicated in the onset of neurological disorders (Zhang et al., 2023). Astrocytes are multifunctional glial cells that play a crucial role in maintaining homeostasis of the central nervous system (CNS), modulating information processing at molecular, cellular and whole network levels, and mediating both beneficial and detrimental effects during neuropathological diseases (Escartin et al., 2021; Lohr, 2023; Verkhratsky & Nedergaard, 2018).

Noradrenergic receptors were initially classified into α and β receptors, the α receptors being further subdivided into α_1_ and α_2_ receptors (Ahlquist, 1948; Lands et al., 1967). A_2_ receptors are known to be coupled to G_i_ and inhibit adenylyl cyclase (AC) activity, whereas β receptors are coupled to G_s_, leading to AC stimulation, and α_1_ receptors are coupled to G_q_, stimulating PLC/IP_3_-dependent Ca^2+^ release from internal stores (Neves et al., 2002; Strosberg, 1993). Astrocytes throughout the brain exhibit Ca^2+^ transients following activation of α_1_ receptors (Wahis & Holt, 2021). However, astrocytic processes in the OB are located in close proximity to noradrenergic fibers and respond to NE application with Ca^2+^ signaling not only via α_1_ but also α_2_ receptors (Fischer et al., 2021). Besides Ca^2+^, cyclic adenosine monophosphate (cAMP) is another ubiquitous second messenger, synthesized from ATP by ACs. cAMP regulates a plethora of cellular functions such as gene transcription and translation. It plays significant roles in neurodevelopment and activity-dependent modifications of neuronal performance (Chen et al., 2022). Astrocytes respond to several neurotransmitters and other signaling molecules with increases in cAMP, including NE (Horvat et al., 2016; Oe et al., 2020), dopamine (von Kalben et al., 2024), adenosine (Theparambil et al., 2024; Wendlandt et al., 2023) and lactate (Vardjan et al., 2018). In astrocytes, cAMP signaling is involved in glucose metabolism (Choi et al., 2012; Hasel et al., 2017; Theparambil et al., 2024; Vardjan et al., 2018), local regulation of blood flow (Vittani et al., 2025), exocytosis (Vardjan & Zorec, 2015), synaptic plasticity (Sitjà-Roqueta et al., 2025) and behavior such as vigilance and memory formation (Oe et al., 2020; Shigetomi & Koizumi, 2023; Zhou et al., 2021; Zorec et al., 2015). To date, ten isoforms of ACs have been characterized in mammals. Nine of them are membrane bound, activated by G_s_ and inhibited by G_i_ (Devasani & Yao, 2022). Three of these AC isoforms (AC1, AC3, and AC8) are Ca^2+^/CaM-stimulated, in addition to their activation by G_s_ (Chen et al., 2022). They could play a central role in the integration between Ca^2+^ and cAMP signaling, but direct evidence for this crosstalk is missing in astrocytes (Sobolczyk & Boczek, 2022). In addition, soluble adenylyl cyclase isoform 10 can be activated by Ca^2+^ and HCO_3-_, while being independent of G_s_ and G_i_ (Li et al., 2024). Ca^2+^-dependent adenylyl cyclases enable functional interaction between both second messenger systems, however, no detailed study of Ca^2+^/cAMP interactions on the single cell level exists in astrocytes.

In the present study, we investigated NE-induced cAMP and Ca^2+^ signals in astrocytes of the mouse OB, which is the first center of olfactory information processing (Rotermund et al., 2019). Although NE-induced cAMP signaling has been extensively studied in various cortical regions and in the hippocampus, noradrenergic cAMP signaling remains underexplored in astrocytes of the OB, despite its significant innervation by projections from the LC (Gao et al., 2016; Nomura et al., 2014; Oe et al., 2020; Shipley et al., 1985). To investigate cAMP dynamics in the olfactory bulb, we expressed the genetically encoded cAMP sensor Flamindo2 (Odaka et al., 2014) in astrocytes and performed confocal cAMP imaging on acute olfactory bulb slices. Ca^2+^ was visualized by GCaMP6s and jRCaMP1a expressed by astrocytes. Our results demonstrate that NE induces cAMP signals in OB astrocytes via α_1_, α_2_, and β receptors, while Ca^2+^ signals were triggered by α_1_ and α_2_ receptors only. Notably, rises in the cAMP concentration via activation of α receptors were dependent on Ca^2+^ release from internal stores such as the ER, whereas cAMP signals induced via β receptors were entirely Ca^2+^-independent. cAMP signals evoked by the α_1_ agonist phenylephrine (PE) were blocked by antagonists of α_1A_ and α_1D_ subtypes and were suppressed by inhibitors of AC1 and AC3. We conclude that NE directly induces an increase in cAMP by activation of β receptors coupled to G_s_, and indirectly by activation of α_1_ and α_2_ receptors leading to Ca^2+^ release from internal stores, thereby stimulating Ca^2+^/CaM-dependent AC1 and AC3.

## Results

### Norepinephrine evokes cAMP signals in astrocytes of the olfactory bulb

To study NE-induced cAMP signaling in astrocytes of the OB we used the genetically encoded cAMP indicator Flamindo2 that consists of circularly permuted Citrine, a mutant of yellow fluorescent protein, fused with the cAMP binding domain of mouse EPAC1 (exchange protein directly activated by cAMP; Fig. 1A, B) (Odaka et al., 2014). We took advantage of AAV-driven Flamindo2 expression controlled by the astrocyte-specific gfaABC1D-promoter (Lee et al., 2008) after retro-bulbar injection and used brain slices including the olfactory bulb 3-6 weeks after virus injection (Fig. 1C-E). We confirmed that Flamindo2 is expressed exclusively in astrocytes by its co-localization with the astrocytic markers GFAP and S100B (Fig. 1F, G). Approximately 50 % of the astrocytes in the glomerular and external plexiform layer of the OB were transduced. Conformational changes in Flamindo2 upon cAMP binding decreases fluorescence (Odaka et al., 2014) (Fig. 1H). Note that subsequent Flamindo2 traces have been inverted (−ΔF) to represent increases in cAMP more intuitively (Fig. 1I; see (Wendlandt et al., 2023)). The cAMP sensor Flamindo2 is pH-sensitive and NE-evoked fluorescence changes could be due to pH changes rather than cAMP changes (Odaka et al., 2014). Therefore, we tested the effect of NE application on intracellular pH in astrocytes loaded with the pH indicator pHrodo Red (fig. s1A). Application of NE did not evoke changes in pH. In contrast, addition of 5 mM NH_4_Cl evoked an acidification that was accompanied by a decrease in Flamindo2 fluorescence, confirming that Flamindo2 fluorescence is pH-sensitive. Our results indicate that NE-induced Flamindo2 fluorescence signals were not mediated by pH changes but reflect cAMP changes (fig. s1B).

**Fig. 1.**
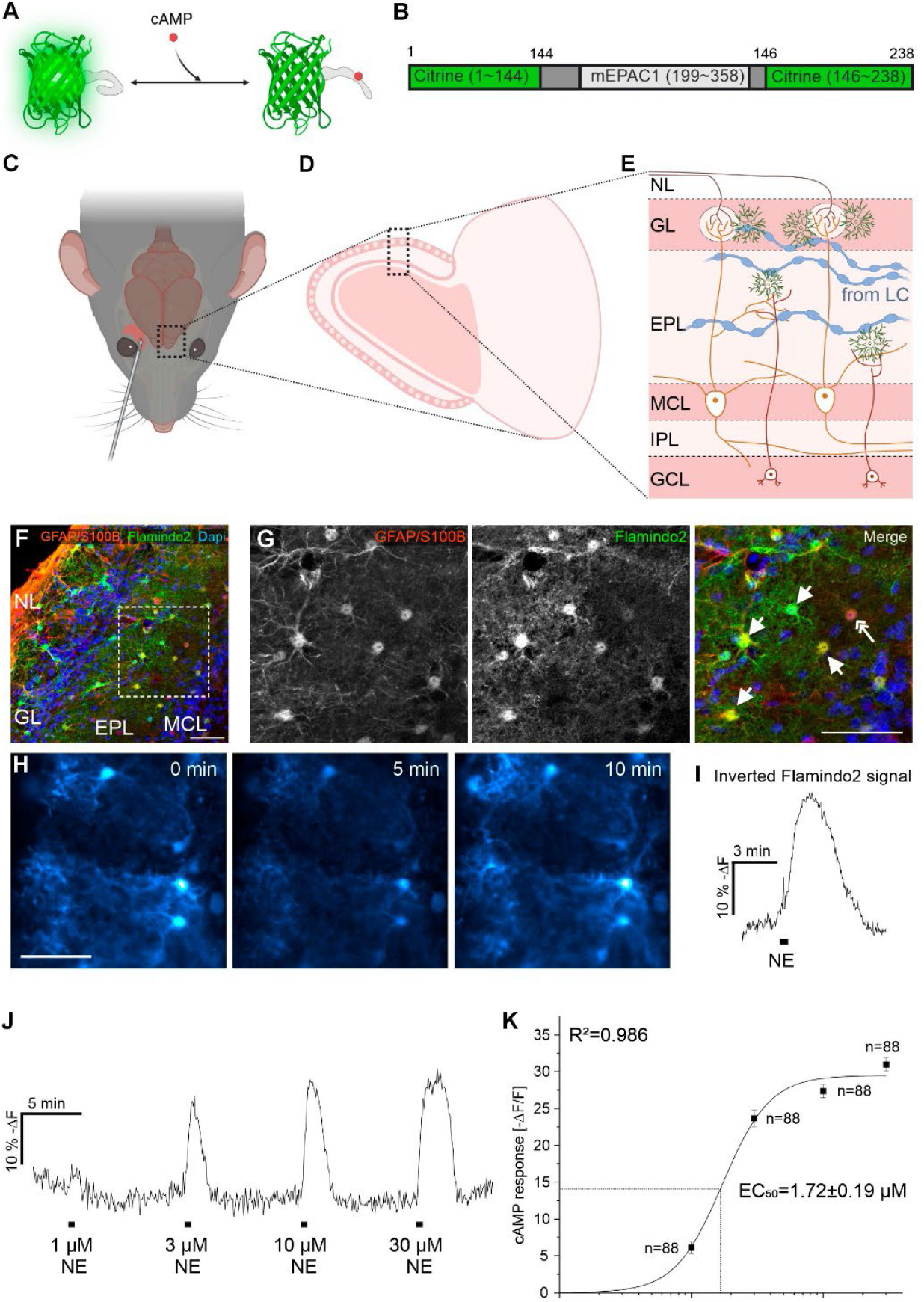
Norepinephrine induces cAMP signals in astrocytes of the olfactory bulb. **(A)** Schematic structure of Flamindo2. Note the decrease in fluorescence upon binding of cAMP. **(B)** Flamindo2 is derived from the yellow fluorescent protein (YFP) mutant, Citrine, and includes the cAMP binding site of mouse EPAC1 (exchange protein directly activated by cAMP) (Odaka et al., 2014). **(C)** Adeno-associated viral vectors (AAVs) carrying the Flamindo2 gene were injected into the retro-bulbar sinus. **(D)** Structure of a sagittal slice of the olfactory bulb. **(E)** Simplified cellular organization of the olfactory bulb: Axons of sensory neurons in the olfactory epithelium enter the nerve layer (NL) and form synapses with mitral cells in the glomerular layer (GL). Dendrites of mitral cells extend through the external plexiform layer (EPL). Cell bodies of mitral cells are located in the mitral cell layer (MCL), their axons proceeding through the internal plexiform layer (IPL) towards the olfactory cortex. The granular cell layer (GCL) contains the cell bodies of granular cells. Centrifugal fibers from the locus coeruleus (LC) containing varicosities innervate the olfactory bulb. Mitral cells are shown in orange color, granular cells in red, and astrocytes in green, centrifugal fibers in blue. **(F)** Immunostaining against Flamindo2 (anti-GFP; green) confirms the expression of Flamindo2 in olfactory bulb astrocytes labeled with anti-GFAP/anti-S100B (red). Nuclei were labeled with Dapi (blue). Scale bar: 50 µm **(G)** Magnified view of astrocytes in the olfactory bulb. Arrows indicating co-expression of Flamindo2 and GFAP/S100B, the double arrow indicates an astrocyte that is Flamindo2-negative. Scale bar: 50 µm. **(H)** Fluorescence of a Flamindo2-expressing olfactory bulb slice prior to (0 min) and during a 30 s application of 10 µM norepinephrine (NE) (5 min). Flamindo2 fluorescence recovered following NE washout (10 min). Scale bar: 50 µm. **(I)** Exemplary Flamindo2 fluorescence trace of a single astrocyte during NE application (10 µM, 30 s). Note the inverted vertical axis (−ΔF) applied to better reflect the changes in astrocytic cAMP concentration. **(J)** Astrocytic cAMP signals evoked by a range of NE concentrations. **(K)** Dose-response-curve of norepinephrine-induced cAMP signals. Data from 3 mice.

To elucidate the dose/response relationship of NE-induced cAMP signals in astrocytes, we applied different concentrations of NE (Fig. 1J). Application of 1 to 30 µM NE led to a dose-dependent increase in cAMP signals with an EC_50_ of 1.72 ± 0.19 µM (Fig. 1K; table s1). All astrocytes expressing Flamindo2 responded to concentrations of 10 µM and higher, and we applied 10 µM NE to induce maximal cAMP responses in OB astrocytes in further experiments

### NE-induced cAMP signals are independent of neuronal activity and mediated via α_1_, α_2_, and β adrenergic receptors

Since NE increases the excitability of neurons in the OB (Linster et al., 2011), we used 0.5 µM tetrodotoxin (TTX) to suppress action potentials in neurons contributing to NE-evoked cAMP signaling in astrocytes. NE was able to induce increases in cAMP when action potentials were blocked with TTX, however, the amplitude of NE-evoked cAMP signals decreased from 37.0 ± 1.3 % −ΔF (n = 92) without TTX to 24.0 ± 0.8 % −ΔF (n = 144) with TTX, indicating some contribution from neurons (fig. s2A-C). To block any residual, TTX-insensitive neuronal effect, we assessed NE-induced cAMP responses in the additional presence of glutamatergic and GABAergic receptor inhibitors using a blocker mix (BM) consisting of TTX (0.5 µM), the NMDA receptor antagonist D-APV (50 µM), the AMPA receptor antagonist NBQX (10 µM), the GABA_A_ receptor antagonist gabazine (5 µM), and the GABA_B_ receptor antagonist CGP 55845 (10 µM). During BM application, the mean amplitude reached 102 ± 3.8 % of the response in TTX alone, which was not significantly different (p = 0.206; Wilcoxon), indicating that in the presence of TTX, NE directly increases astrocytic cAMP without a significant contribution from neuronal transmitter release (Fig. 2A)

**Fig. 2.**
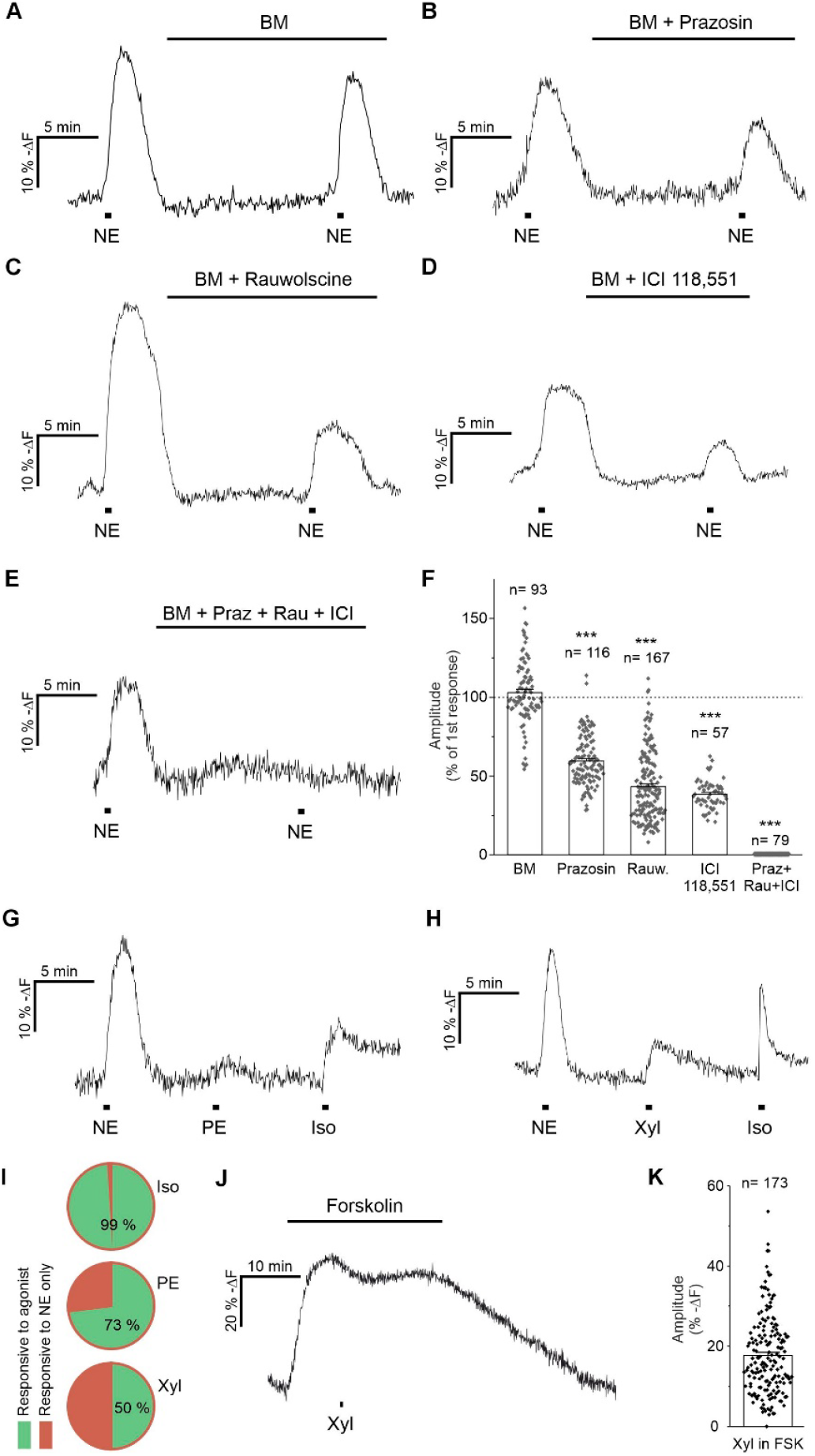
Norepinephrine-induced cAMP signals are independent of neuronal activity and mediated by α_1_, α_2_ and β adrenergic receptors. **(A)** Norepinephrine (NE)-induced cAMP signals in the presence of tetrodotoxin (TTX; first application) and TTX combined with glutamatergic and GABAergic inhibition in a blocker mix (BM) containing: 50 µM DAPV (NMDA receptors), 10 µM NBQX (AMPA receptors), 5 µM Gabazine (GABA_A_ receptors), 10 µM CGP 55845 (GABA_B_ receptors). **(B-E)** Inhibition of NE-induced cAMP signals by the noradrenergic antagonists **(B)** prazosin (α_1_; 10 µM), **(C)** rauwolscine (α_2_; 0.5 µM), **(D)** ICI 118,551 (β; 15 µM), and **(E)** the combination of all three antagonists. **(F)** Quantification of amplitudes of cAMP transients normalized to the amplitude of the first NE application. ***; p < 0.001, Kruskal Wallis ANOVA and Dunńs post hoc test. Data from 3 mice (prazosin), 3 mice (rauwolscine), 3 mice (ICI118,551) and 3 mice (Praz+Rau+ICI). **(G)** cAMP signals evoked by application of 10 µM NE, 100 µM phenylephrine (PE; α_1_ agonist), 100 µM isoprenaline (Iso; β agonist), and **(H)** 10 µM NE, 80 µM xylazine (Xyl; α_2_ agonist) and 100 µM Iso. **(I)** Fraction of astrocytes responding to Iso (total of 87 cells from 3 mice), PE (total of 30 cells from 3 mice) and Xyl (total of 66 cells from 3 mice). **(J)** Inhibitory effect of Xyl during elevation of cAMP levels upon adenylyl cyclase stimulation by 3 µM forskolin. **(K)** Amplitudes of xylazine-evoked decreases in cAMP during application of forskolin. Data from 4 mice.

We tested the involvement of α_1_, α_2_, and β adrenergic receptor subtypes by assessing the effect of subtype-specific antagonists on NE-induced cAMP signaling in astrocytes in the presence of TTX and glutamatergic/GABAergic antagonists. The antagonists prazosin (10 µM, α_1_ receptor) and rauwolscine (0.5 µM, α_2_ receptor) significantly decreased NE-evoked astrocytic cAMP responses to 60 ± 1.5 % (n = 116; Fig. 2B) and 43 ± 1.7 % (n = 167; Fig. 2C) of their respective controls. The β receptor antagonist ICI 118,551 (15 µM) reduced the NE-induced cAMP responses to 38 ± 1.2 % of the control (n = 57; Fig. 2D), whereas a combination of all three antagonists completely blocked NE-evoked increases in cAMP (n = 79; Fig. 2E). These results, summarized in Fig. 2F, suggest that all three noradrenergic receptor subtypes contribute to astrocytic cAMP. We next used subtype-specific agonists (in the presence of 0.5 µM TTX) to verify the results. 100 µM phenylephrine (PE; α_1_ receptor agonist), 80 µM xylazine (α_2_ receptor agonist) and 100 µM isoprenaline (β receptor agonist) increased cAMP in OB astrocytes (Fig. 2G, H). Isoprenaline induced the largest cAMP signals with an amplitude of 22.3 ± 0.7 % −ΔF (n = 86 out of 87 cells), which were significantly larger than those induced by PE (5.6 ± 0.4 % −ΔF, n = 22 out of 30 cells; p < 0.001) and xylazine (10.6 ± 0.7 % −ΔF, n = 33 out of 66 cells; p < 0.001) (fig. s3). Whereas 99 % of the astrocytes responding to NE also increased cAMP in response to isoprenaline, only 73 % and 48 % of NE-responsive astrocytes responded to PE and xylazine, respectively (Fig. 2I). Since α_2_ receptors are known to couple to G_i_ and inhibit cAMP production, the cAMP increase mediated by the α_2_ receptor agonist xylazine was surprising. Interestingly, xylazine was also able to decrease 3 µM forskolin-induced cAMP production, indicating that α_2_ receptors in OB astrocytes couple to the G_i_ adenylyl cyclase-inhibiting pathway in addition to stimulating cAMP production (−17.7 ± 0.7 % −ΔF from forskolin peak, n = 173, Fig. 2J, K).

### NE-induced cAMP signals partly depend on Ca^2+^ signals

Our results indicate that in OB astrocytes, not only β-adrenergic receptors but also α_1_ and α_2_ receptors trigger cAMP production. Previously, we showed that α_1_ and α_2_ receptors increase Ca^2+^ in OB astrocytes (Fischer et al., 2021), raising the question whether NE-evoked Ca^2+^ activates ACs to increase cAMP. To investigate Ca^2+^ and cAMP signals simultaneously, we combined the red fluorescent Ca^2+^ indicator jRCaMP1a with Flamindo2 (Fig. 3A, B). Since repetitive application of NE leads to diminishing Ca^2+^ transients (rundown) (Fischer et al., 2021), we first characterized the rundown of NE-induced cAMP signals (Fig. 3C). Initial applications of NE led to cAMP and Ca^2+^ signals with amplitudes of 24.0 ± 0.8 % −ΔF for cAMP (n=144) and 31.5 ± 2.3 % ΔF for Ca^2+^ (n = 36; Fig. 3C). Second applications resulted in cAMP and Ca^2+^ transients that were 92 ± 1.9 % of the first response for cAMP (n = 144; p < 0.001) and 78 ± 2.2 % of the first response for Ca^2+^ (n = 36; p < 0.001). NE-evoked Ca^2+^ signals consisted of two components; a fast, transient Ca^2+^ rise reflecting Ca^2+^ release from intracellular stores, and a sustained plateau phase reflecting activation of store-operated Ca^2+^ entry upon store depletion, the latter being best described by the area under the curve (Fischer et al., 2021). Therefore, we additionally calculated the area under the curve of the Ca^2+^ signals, which was reduced to 74 ± 3.6 % of the first response upon a second application of NE (n = 36; Fig. 3C, G).

**Fig. 3.**
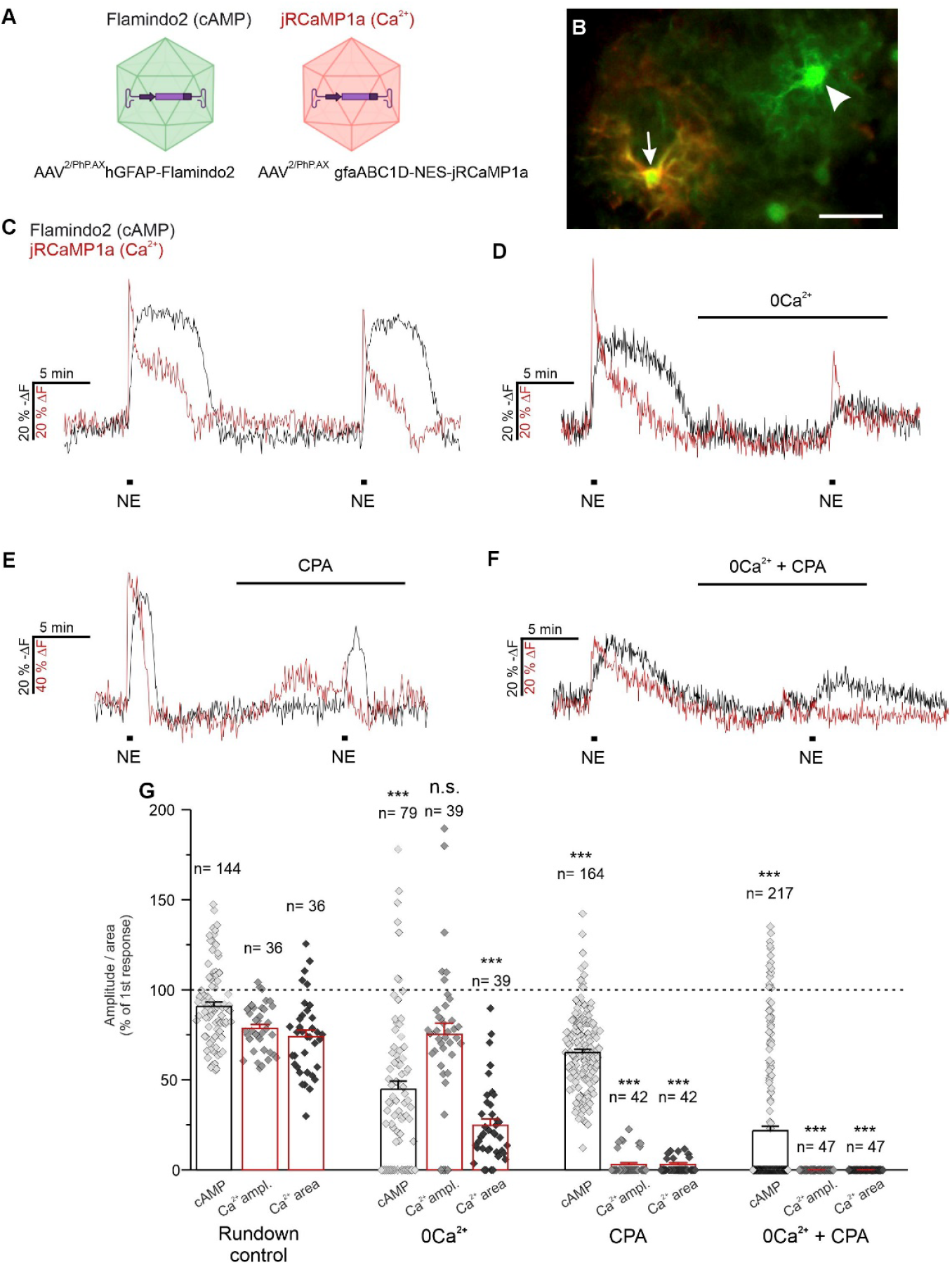
Norepinephrine-induced cAMP signals are partly Ca^2+^-dependent. **(A)** Astrocytes were co-transduced with AAVs carrying Flamindo2 or jRCaMP1a for simultaneous cAMP and Ca^2+^ measurements. **(B)** Astrocytes in an acute brain slice co-transduced with Flamindo2 and jRCaMP1a (arrow) and Flamindo2 alone (arrowhead). Scale bar: 30 µm. **(C)** cAMP (black) and Ca^2+^ signals (red) evoked by repetitive application of 10 µM norepinephrine (NE) to determine rundown effects. **(D)** Effect of calcium-free ACSF (0Ca^2+^), **(E)** Ca^2+^ store depletion by 20 µM cyclopiazonic acid (CPA), and **(F)** the combination of calcium-free ACSF and CPA on NE-evoked responses. **(G)** Quantification of cAMP and Ca^2+^ transients, compared to the corresponding response of the rundown experiment. For Ca^2+^ responses both the amplitude as well as the area under the curve were analysed. Note that most data points for Ca^2+^ amplitude and Ca^2+^ area with CPA and 0Ca^2+^/CPA have a value of zero and lie on the baseline. ***p < 0.001, n.s.: not significant, Kruskal Wallis ANOVA and Dunńs post hoc test. Some experiments were performed on mice only transduced with Flamindo2. Rundown control: 3 mice Flamindo2, 3 mice Flamindo2 + jRCaMP1a; 0Ca^2+^: 1 mouse Flamindo2, 3 mice Flamindo2 + jRCaMP1a; CPA: 3 mice Flamindo2, 3 mice Flamindo2 + jRCaMP1a; 0Ca^2+^ + CPA: 3 mice Flamindo2, 3 mice Flamindo2 + jRCaMP1a.

We used the second application of the rundown experiment in Fig. 3C as the control group and compared these Ca^2+^ and cAMP transients to the responses after pharmacological manipulation in the following experiments. To test whether Ca^2+^ is essential for cAMP signaling, we either omited Ca^2+^ in the ACSF to prevent Ca^2+^ influx or inhibited the sarco-/endoplasmic reticulum Ca^2+^ ATPases (SERCA) with 20 µM cyclopiazonic acid (CPA) to deplete intracellular Ca^2+^ stores and suppress intracellular Ca^2+^ release. Ca^2+^ store depletion was visible by a moderate increase in cytosolic Ca^2+^ which was often accompanied by a slow increase in cAMP (fig. s4A). After application of Ca^2+^-free ACSF for ten minutes, the peak Ca^2+^ response to NE was unaffected (75 ± 6.3 % of the first response. n = 39; p > 0.05 vs rundown control), while the area under the curve was significantly reduced to 25 ± 3.4 % of the first response (n = 39; p < 0.001), confirming that the plateau phase but not the fast peak depends on Ca^2+^ influx (Fig. 3D, G).

Depleting extracellular Ca^2+^ significantly reduced cAMP signals to 45 ± 4.6 % of the first response (n = 79; p < 0.001). Depleting intracellular Ca^2+^ stores by CPA almost entirely suppressed NE-evoked Ca^2+^ transients (n = 42) and significantly reduced cAMP signals to 65 ± 1.7 % of the first response (n = 164; p < 0.001, Fig. 3E, G). When Ca^2+^-free ACSF and CPA were combined, NE-induced Ca^2+^ signals were completely eliminated (n = 47; Fig. 3F, G) whereas cAMP signals with an amplitude of 22 ± 2.6 % of the first response persisted (n = 217; p < 0.001). Our results show that mechanisms whereby NE increases cAMP in OB astrocytes consists of a Ca^2+^-dependent and a Ca^2+^-independent component.

β receptors are known to directly stimulate adenylyl cyclases by G_s_, suggesting that the Ca^2+^-independent pathway is stimulated by β receptors. Indeed, the β receptor agonist isoprenaline failed to induce Ca^2+^ signals (fig. s4B, C). cAMP signals evoked by isoprenaline were not affected when Ca^2+^ stores were emptied by CPA, confirming that cAMP production by β receptors is Ca^2+^-independent (fig. s4D-F).

### PE-induced cAMP signals are Ca^2+^-dependent

Activation of α_1_ and α_2_ receptors increases Ca^2+^ in OB astrocytes (Fischer et al., 2021) and canonically they are not linked to G_s_ proteins, suggesting that the Ca^2+^-dependent increases in cAMP in our study is mediated by α_1_ and α_2_ receptors. Ca^2+^-dependent cAMP signaling in OB astrocytes is best studied by selective activation of α_1_ receptors by, e.g., PE, since α_2_ receptors both increase cAMP and decrease adenylyl cyclase activity in OB astrocytes. PE at concentrations between 1 and 300 µM led to dose-dependent cAMP signals with amplitudes ranging from 3.7 ± 0.5 % −ΔF (1 µM PE) to 8.7 ± 0.5 % −ΔF (300 µM PE; n = 29–112; Fig. 4A; table s2). We calculated the EC_50_ to be 1.26 ± 0.33 µM (Fig. 4B). For further experiments, we used 10 µM PE to elicit robust sub-saturating cAMP signals. We applied PE twice in rundown experiments with intervals of 10 min (RD short) and 30 min (RD long) between the cAMP signals and compared these rundowns to any further experiment (fig. s5A-D). A second application of PE evoked cAMP responses in RD short of 84 ± 3.7 % of the first application (n = 29) and in RD long of 79 ± 5.7 % of the first application (n = 40).

**Fig. 4.**
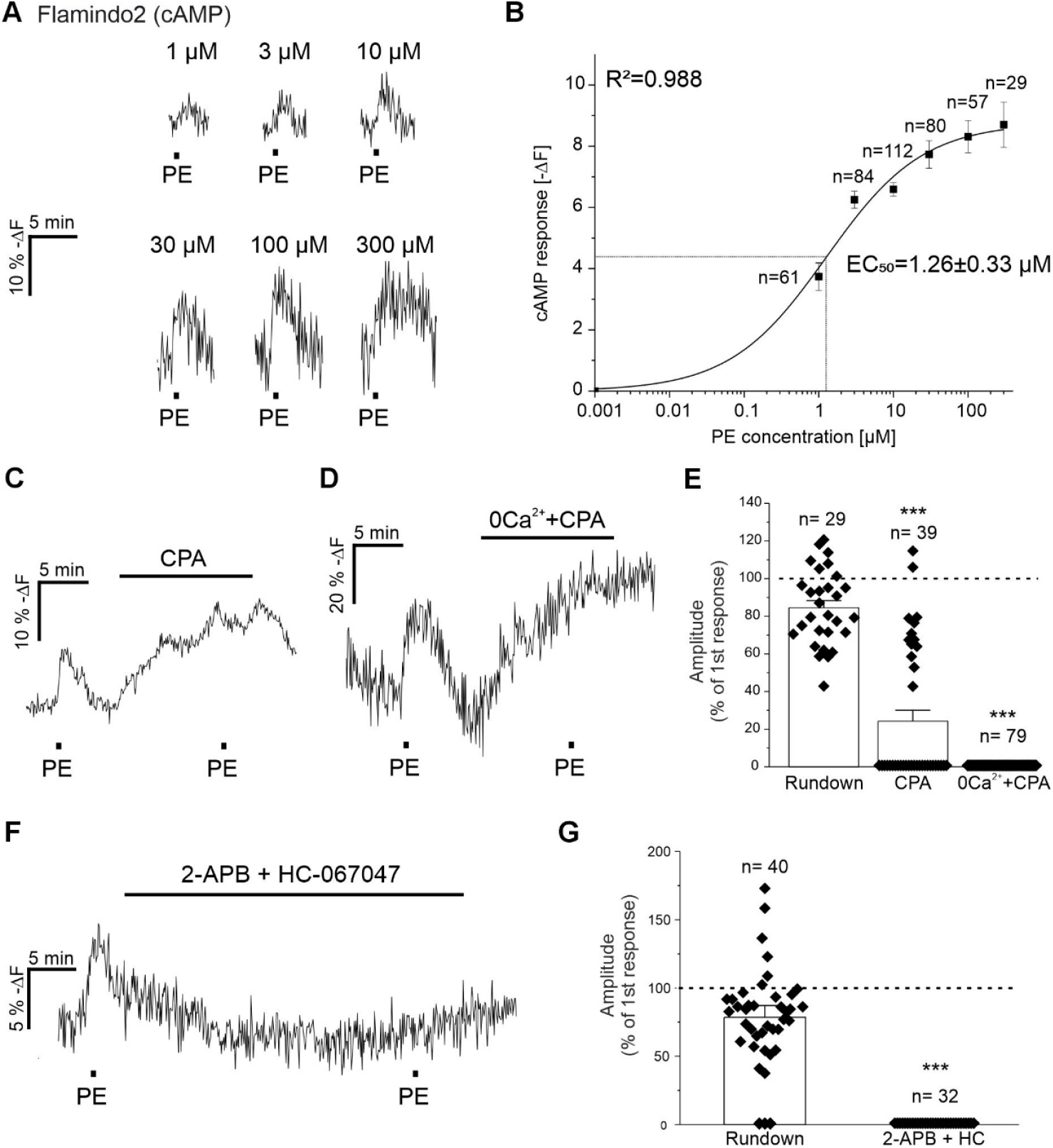
α_1_ receptor agonist phenylephrine induces Ca^2+^-dependent cAMP signals. **(A)** cAMP signals in olfactory bulb astrocytes evoked by phenylephrine (PE) at concentrations ranging from 1 µM to 300 µM. **(B)** Dose-response curve of PE-induced cAMP signals. Experiments from 6 mice. **(C)** PE-induced cAMP signals in the presence of 20 µM cyclopiazonic acid (CPA) and **(D)** the combination of calcium-free ACSF (0Ca^2+^) and CPA. **(E)** CPA and CPA/0Ca^2+^ significantly reduced the amplitude of PE-evoked cAMP signals when compared to the corresponding rundown experiment (***; p < 0.001, Kruskal Wallis ANOVA and Dunńs post hoc test). 3 mice for rundown experiments, 3 mice for CPA, 4 mice for 0Ca^2+^+CPA. **(F)** PE-induced cAMP signals were suppressed in the presence of the IP_3_ receptor inhibitor 2-APB (100 µM) and the TRPV4 inhibitor HC-067047 (1 µM). **(G)** Quantification of the effect of 2-APB+HC-067047. ***; p < 0.001, Mann-Whitney U-Test; 7 mice for rundown long, 4 mice for 2-APB+HC.

To investigate the Ca^2+^ dependency of PE-induced cAMP signals, we blocked SERCA pumps with 20 µM CPA to deplete Ca^2+^ stores (Fig. 4C). After depletion of Ca^2+^ stores by incubation of CPA for 10 min, PE-induced cAMP signals were significantly smaller compared to the corresponding rundown (RD short; p < 0.001) and averaged 24 ± 5.8 % (n = 39) of the first response, while PE-induced cAMP signals were entirely suppressed in the presence of CPA and Ca^2+^-free ACSF (0Ca^2+^/CPA; Fig. 4C-E). Forskolin-induced cAMP signals, in contrast, were not affected by CPA, showing that CPA does not directly inhibit cAMP production by, e.g., blocking adenylyl cyclases (Fig. s6A-C). To confirm the involvement of intracellular Ca^2+^ release, we inhibited IP_3_ receptors with 100 µM 2-APB, which blocks both IP_3_ receptors and store-operated Ca^2+^ entry (Singaravelu et al., 2006; Stavermann et al., 2012). As 2-APB at concentrations above 50 µM also stimulates TRPV4 channels and Ca^2+^ oscillations in OB astrocytes (Doengi et al., 2009), we co-applied 1 µM of the selective TRPV4 antagonist HC-067047 with 2-APB for 30 min (Fischer et al., 2021). The PE-induced increase in cAMP vanished completely after blocking IP_3_ receptors, indicating that Ca^2+^ release mediated by IP_3_ receptors is required for cAMP production evoked by α_1_ receptors (n = 32; Fig. 4F, G). In conclusion, our results demonstrate that PE-induced Ca^2+^ signals depend on the PLC/IP_3_ pathway as well as Ca^2+^ release from the endoplasmic reticulum and trigger increases in cAMP.

### α_1A_ and α_1D_ receptor subtypes increase Ca^2+^ and cAMP

The α_1_ receptors are classified into the subtypes α_1A_, α_1B_, and α_1D_ (Bylund et al., 1994; Hieble et al., 1995). For further investigation of Ca^2+^ and cAMP signals induced by PE-application, we used specific antagonists for the different α_1_ receptor subtypes. Due to the relatively low co-expression rate of combined jRCaMP1.0 and Flamindo2 AAVs in the previous experiments, we performed the following experiments for Ca^2+^ and cAMP imaging separately, using GLAST-CreERT2 x GCaMP6s^flox^ mice for Ca^2+^ and AAV carrying Flamindo2 for cAMP (Fig. 5A, B).

**Fig. 5.**
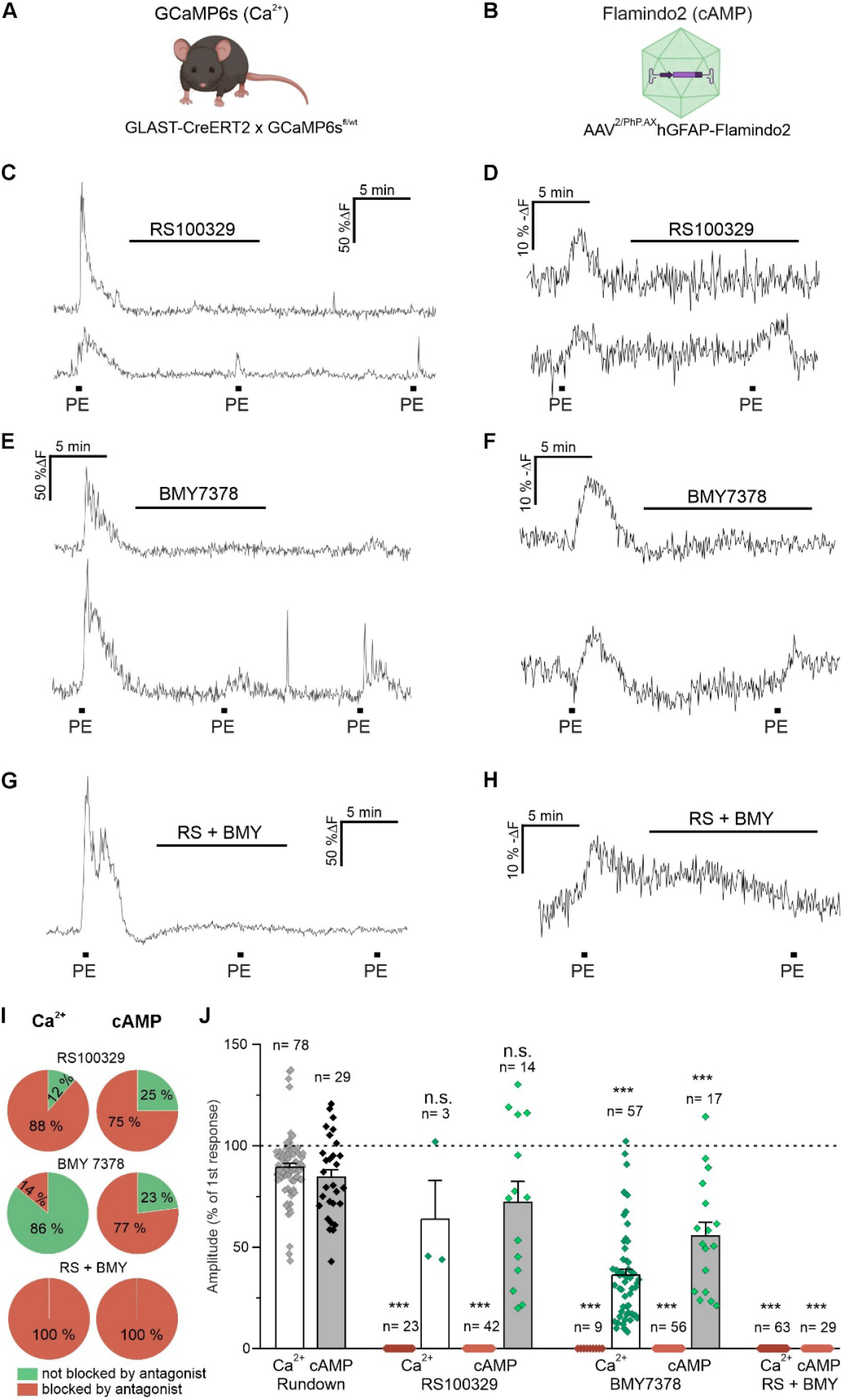
Phenylephrine induces Ca^2+^ and cAMP signals via α_1A_ and α_1D_ receptor subtypes. **(A)** To visualize Ca^2+^ signaling in olfactory bulb astrocytes, GCaMP6s was expressed under control of the promoter of the astrocyte-specific glutamate-aspartate transporter (Glast). **(B)** Flamindo2 was introduced into wild type mice by AAV to visualize cAMP signaling. **(C)** Effect of RS100329 (α_1A_; 1 µM) on phenylephrine (PE)-induced Ca^2+^ signals and **(D)** cAMP signals. RS100329 entirely inhibited Ca^2+^ and cAMP signals in a subpopulation of astrocytes (upper traces), while Ca^2+^ and cAMP signals remained in other astrocytes (lower traces). **(E)** BMY7378 (α_1D_; 1 µM) blocked Ca^2+^ signals and **(F)** cAMP signals in some astrocytes (upper traces) and reduced Ca^2+^ and cAMP signals in the remaining astrocytes (lower traces). **(G)** The combination of RS100329 and BMY entirely inhibited PE-induced Ca^2+^ and **(H)** cAMP signals. **(I)** Fraction of responsive vs. non-responsive astrocytes in the presence of RS100329, BMY7378 and the combination of both. **(J)** Analysis of PE-induced Ca^2+^ and cAMP signals. Note that in the presence of RS100329 as well as BMY7378 two subpopulations of astrocytes could be identified, one with Ca^2+^ and cAMP transients entirely blocked by the antagonists and one with residual Ca^2+^ and cAMP transients in the presence of the antagonists. ***p < 0.001, n.s.= not significant; Kruskal Wallis ANOVA and Dunńs post hoc test. Ca^2+^ imaging: 4 mice for rundown, 3 mice for RS100329, 3 mice for BMY7378, 3 mice for RS100329 + BMY7378. cAMP imaging: 3 mice for rundown, 4 mice for RS100329, 3 mice for BMY7378, 4 mice for RS100329 + BMY7378.

Application of PE evoked Ca^2+^ transients in 95 % of all astrocytes (n = 73 out of 77), i.e. astrocytes that responded to application of NE, indicating that the majority of OB astrocytes express α_1_ receptors linked to Ca^2+^ signaling (fig. s7A). PE-evoked cAMP signals, in contrast, occurred in 53 % of astrocytes only (n = 71 out of 133) in this set of experiments, suggesting that in some cells, α_1_ receptors are not linked to cAMP production (fig. s7B, C). To study the contribution of α_1A_ receptors, we used the α_1A_-specific antagonist RS100329, which had a bimodal effect on both Ca^2+^ and cAMP signals. 1 µM RS100329 entirely blocked PE-induced Ca^2+^ signals in 88 % of astrocytes (n = 23 out of 26), while Ca^2+^ signals were only weakly reduced in the remaining 12 % of the astrocytes and, on average, did not differ significantly from the rundown experiment (p > 0.05; n = 3 out of 26; Fig. 5 C, I and fig. s8A, B).

In addition, RS100329 entirely blocked PE-induced cAMP signals in 75 % of the astrocytes (n = 42 out of 56) and resulted in PE-induced cAMP signals that did not differ significantly from the rundown experiment in the remaining 25 % of astrocytes (n = 14 out of 56; Fig. 5D, I). The α_1D_ receptor antagonist BMY7378 also had a bimodal effect. 1 µM BMY7378 completely blocked PE-induced Ca^2+^ signals in 14 % of the cells (n = 9 out of 66) and reduced Ca^2+^ signals to 36 ± 3.0 % of the initial amplitude in 86 % of the cells (p < 0.001; n = 57 out of 66; Fig. 5E, I). BMY7378 blocked PE-induced cAMP signals in 70 % of the cells (n = 56 out of 73) and reduced cAMP signals to 55 ± 6.9 % of the initial amplitude in the remaining 30 % of the cells (p = 0.171; n = 17 out of 73; Fig. 5F, I). Combination of both blockers completely abolished both PE-induced Ca^2+^ (n = 63; Fig. 5G, I) and cAMP signals (n = 29; Fig. 5H, I). The complete inhibition of the second messengers by combination of α_1A_ and α_1D_ antagonists indicates that α_1B_ does not play a significant role in olfactory bulb astrocytes. In summary, our results show that α_1_ receptors, including α_1A_ and α_1D_, induce Ca^2+^ signaling in virtually all astrocytes of the olfactory bulb, while cAMP signals are evoked in a subpopulation of astrocytes (Fig. 5J).

### Ca^2+^-induced cAMP signals are mediated by AC1 and AC3

Our results show that an increase in Ca^2+^ can lead to an increase in cAMP in OB astrocytes, suggesting the involvement of Ca^2+^-activated adenylyl cyclases. Three out of the ten AC isoforms can be activated by Ca^2+^/CaM: AC1, AC3 and AC8 (Devasani & Yao, 2022). We tested the presence and co-localization with astrocytes of AC1, AC3 and AC8 using antibody staining. We visualized astrocytes of the olfactory bulb with a combination of anti-GFAP and anti-S100B antibodies. Co-staining of astrocytes and AC1 (Fig. 6A, B) and AC3 (Fig. 6C, D), respectively, demonstrated co-localization in the olfactory bulb. For AC8, in contrast, no clear co-localization with astrocytes was found (Fig. 6E, F).

**Fig. 6.**
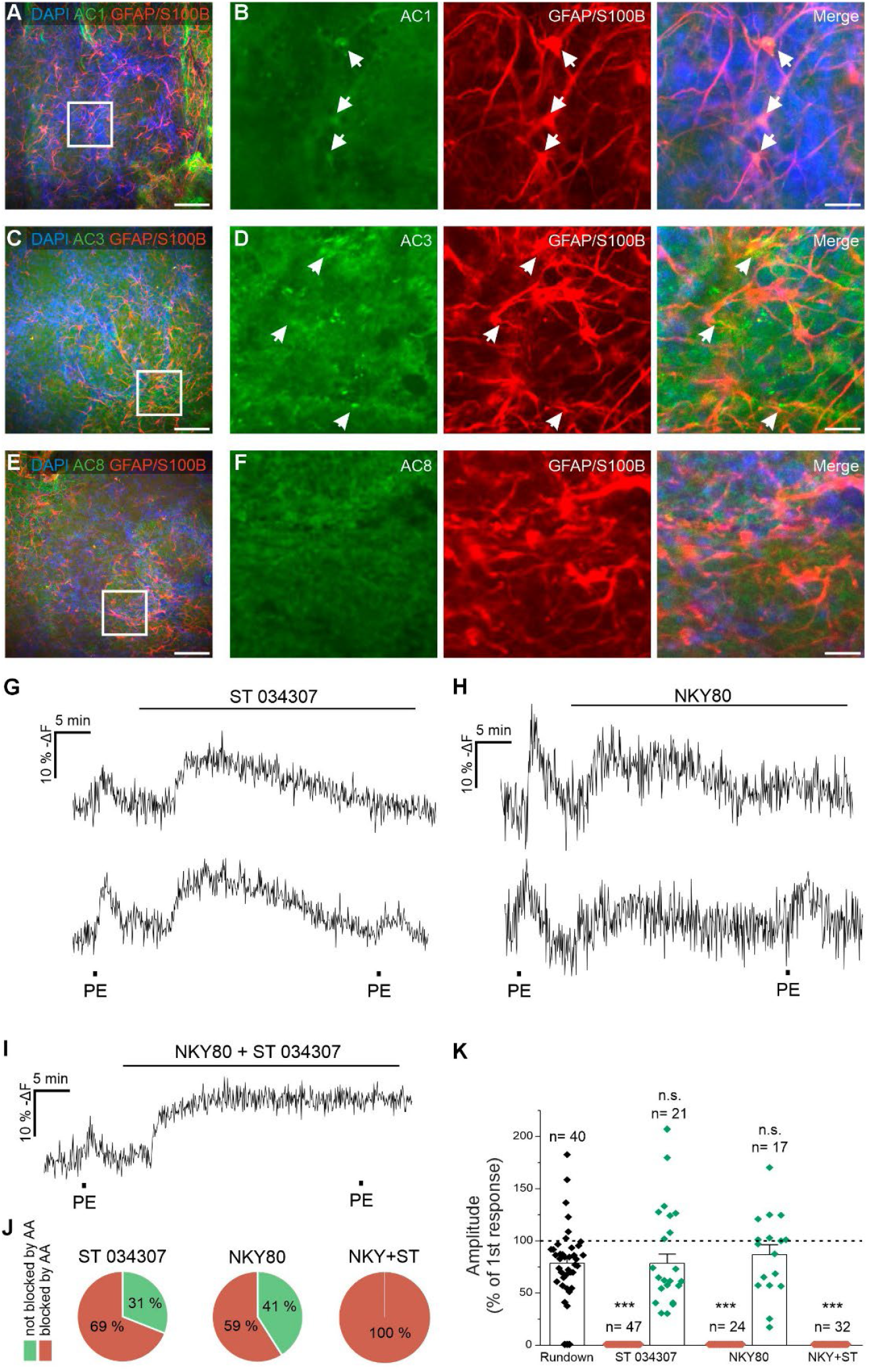
Phenylephrine-induced cAMP signals are mediated via adenylyl cyclase subtypes 1 and 3. **(A)** Anti-GFAP/S100B immuno-staining of astrocytes (red) and anti-adenylyl cyclase 1 immunostaining (AC1; green). Nuclei are stained with Dapi (blue). **(B)** Magnified view, with arrows pointing to astrocytes co-stained with GFAP/S100B and AC1 antibodies. **(C)** Immunostaining of astrocytes (red) and adenylyl cyclase 3 (AC3, green). **(D)** Magnified image, depicting astrocytes co-stained with GFAP/S100B and AC3 antibodies (arrows). **(E)** Immunostaining of astrocytes (red) and adenylyl cyclase 8 (AC8, green). **(F)** In the agnified view, no co-localization of GFAP/S100B and AC8 immunostaining was detected. Scale bars: A, C, E; 50 µM. B, D, F; 10 µM **(G)** Effect of the AC1 blocker ST 034307 (10 µM) on phenylephrine (PE)-induced cAMP signals. ST 034307 entirely blocked cAMP transients in a subpopulation of astrocytes (upper trace), but had no significant effect on the cAMP transients in the remaining astrocytes (lower trace). **(H)** Effect of the AC3-blocker NKY80 (500 µM) on PE-induced cAMP signals. In some astrocytes, NKY80 entirely inhibited cAMP transients (upper trace), while in the remaining astrocytes it had no significant effect on cAMP transients (lower trace). **(I)** Combination of ST 034307 and NKY80 completely blocked PE-induced cAMP signals. **(J)**. Fraction of astrocytes responding to NE vs. not responding to NE in the presence of ST 034307, NKY80 and the combination of ST 034307 and NKY80. **(K)** Analysis of the effects of the adenylyl cyclase inhibitors on PE-evoked cAMP signals. ***p < 0.001, n.s.= not significant; Kruskal Wallis ANOVA and Dunńs post hoc test. 7 mice for rundown, 5 mice for ST 034307, 5 mice for NKY80, 3 mice for ST 034307 + NKY80.

To test the contribution of Ca^2+^-activated AC1 and AC3 isoforms to cAMP signaling in OB astrocytes, we used the AC1-specific antagonist ST 034307, as well as NKY80, an antagonist of several adenylyl cyclases including AC3 (Brand et al., 2013; Brust et al., 2017). Blockage of AC1 with 10 µM ST 034307 completely inhibited PE-induced cAMP responses in 69 % of the astrocytes (n=47 out of 68), while in the remaining 31 % of the astrocytes, the response did not differ significantly from the rundown experiment (n=21 out 68; p=1; Fig. 6G, J). In addition, NKY80 (500 µM) completely blocked cAMP signals in 59 % of the cells (n=24 out of 41), whereas in the remaining 41 % of the cells, the cAMP signal was not statistically different from the rundown experiment (n=17 out of 41; p=1; Fig. 6H, J). The combined blockage of AC1 and AC3 led to a total inhibition of the PE-induced cAMP signal (n=32; Fig. 6I-K). In our experiments, application of the AC inhibitors itself led to an increase in the cAMP concentration, which might result from them being a positive modulator of Ca^2+^-independent AC2 (Brust et al., 2017). In conclusion, our results suggest that stimulation of α_1_ receptors by PE induces Ca^2+^-dependent cAMP signals in olfactory bulb astrocytes via activation of AC1 and AC3.

## Discussion

In this study, we investigated the mechanisms by which NE evokes cAMP signaling in astrocytes of the glomerular and external plexiform layers of the OB. Our data demonstrate that NE induces cAMP generation via all three adrenergic receptor classes, i.e., α_1_, α_2_, and β receptors. While β receptors evoke cAMP signals in all OB astrocytes in a Ca^2+^-independent manner by direct stimulation of G_s_-proteins, α_1_-mediated cAMP responses as well as α_2_-mediated cAMP responses occur in a subpopulation of astrocytes and require intracellular Ca^2+^ rises to activate Ca^2+^-sensitive adenylate cyclases AC1 and AC3. These findings highlight a complex, receptor- and Ca^2+^-dependent regulation of cAMP signaling that has not been described in astrocytes before.

### NE induces cAMP signals in astrocytes of the olfactory bulb via α_1_, α_2_, and β receptors

We determined an EC_50_ of approximately 2 µM for NE-induced cAMP responses in OB astrocytes and used 10 µM for all further experiments, a concentration also applied in previous studies addressing Ca^2+^ signaling in olfactory bulb astrocytes and excitability of olfactory bulb neurons (Fischer et al., 2021; Zimnik et al., 2013). Importantly, suppression of neuronal activity and synaptic transmission only partly reduced NE-induced cAMP signals, suggesting that the remaining effect is a direct consequence of NE acting on astrocytes, independent of neuronal input. Astrocytes are known to participate in the “glutamate amplifies noradrenergic effects” (GANE) mechanism by releasing glutamate that triggers neuronal activity and hence further NE release (Mather et al., 2016; Wahis & Holt, 2021). As inhibiting neuronal activity decreased NE-evoked rises in cAMP, such a feedback loop also appears to exist in the olfactory bulb.

We found that agonists for α_1_, α_2_, and β receptors each induced cAMP elevations, indicating that all three receptor types contribute to cAMP signaling under resting conditions, which is in contrast to, e.g., hippocampal astrocytes in which β receptors increase cAMP, and only α_1_ receptors increase Ca^2+^ (Oe et al., 2020). Given NE’s binding affinities at these receptors (α_2_ > α_1_ > β; (Ramos & Arnsten, 2007)), even low NE levels could recruit α_2_ and α_1_ receptors, while higher concentrations are needed to activate β receptors.

An unexpected result was the increase in cAMP evoked by α_2_ receptor activation. Astrocytes in the cortex show opposing regulation of cAMP by α_2_ and β receptors, starting from astrocytic cAMP resting state (Rosenberg et al., 2023), while in our experiments the α_2_ agonist xylazine decreased cAMP only when ACs were pre-activated with forskolin. Hence, α_2_ receptors bimodally regulate cAMP signals depending on the activation state of the astrocyte, increasing cAMP at resting cAMP levels and decreasing cAMP when ACs are already active and cAMP levels are elevated.

### NE-induced and PE-induced cAMP signals differ in their Ca^2+^ dependency

Previous work has shown NE-induced Ca^2+^ transients in astrocytes via α_1_ and α_2_ receptors (Fischer et al., 2021), and we now extend these findings by demonstrating that NE-induced Ca^2+^ signals increase cAMP. SERCA inhibition with CPA reduced NE-induced cAMP signals by approximately 35 %. The residual cAMP responses were Ca^2+^-independent and persisted even in Ca^2+^-free extracellular solution, suggesting a direct activation of ACs by G_s_ proteins as described for signaling mediated by β receptors. In contrast to NE-evoked cAMP signals, PE-induced α_1_-mediated cAMP signals were entirely dependent on Ca^2+^ signaling. Although CPA alone did not fully block these responses, the combination of CPA with Ca^2+^-free ACSF as well as IP_3_ receptor antagonism entirely abolished cAMP generation. These results suggest that the initial ER-derived Ca^2+^ release and subsequent store-operated Ca^2+^ entry (SOCE) are essential for PE-induced AC activation. Interestingly, while activation of α_1_ receptors have previously been associated primarily with Ca^2+^ signaling in astrocytes of various brain regions, as shown *in vivo* in the cerebral cortex (Oe et al., 2020), cAMP responses to α_1_ receptor activation have been reported in peripheral systems such as the uterine smooth muscle cells (Chen et al., 2018). Our data show that OB astrocytes also have α_1_ receptor coupling to increases in cAMP via Ca^2+^-dependent AC1 and AC3. This highlights functional diversity in α_1_-dependent signaling across brain regions and the linkage of the two second messenger pathways in astrocytes.

### PE-induced cAMP signals require α _1A_ and α _1D_ receptors and subsequent activation of AC1 and AC3

We identified both α _1A_ and α _1D_ noradrenergic receptors as key mediators of PE-induced Ca^2+^ and cAMP signals in OB astrocytes. However, there is a discrepancy between Ca^2+^ and cAMP signaling regarding the cell fractions that are sensitive to inhibition of α _1A_ versus α _1D_ receptors (Fig. 5I). Blocking α _1A_ receptors completely inhibited PE-induced Ca^2+^ signals in 88 % of cells, whereas blocking α _1D_ receptors entirely blocked PE-induced Ca^2+^ signals in only 14 % of the cells, suggesting that both astrocyte populations complete each other, with α _1A_ being sufficient to elicit a Ca^2+^ rise in the majority of astrocytes. PE-induced cAMP signals were entirely blocked by inhibition of α _1A_ receptors in 75 % of the cells, a fraction similar to that found in Ca^2+^ imaging experiments. Inhibition of α _1D_ receptors by BMY 7378 suppressed cAMP signals in about 80 % of the astrocytes, a cell fraction much larger compared to the fraction of astrocytes in which Ca^2+^ signaling was entirely inhibited by BMY 7378. However, Ca^2+^ signals in astrocytes that were not entirely blocked by BMY 7378 were strongly attenuated, indicating that α _1D_ receptors, while not essential to evoke Ca^2+^ signals, augment Ca^2+^ signals induced by α_1A_ receptors. Consequently, attenuated PE-evoked Ca^2+^ signals in the presence of BMY 7378 were insufficient to stimulate Ca^2+^-dependent ACs and failed to elicit measurable cAMP signals in the majority of astrocytes. Our data identified AC1 and AC3 as the primary ACs responsible for PE-induced cAMP production using the selective AC1 inhibitor ST034307 and the non-specific AC inhibitor NKY80. NKY80 is known to mainly inhibit AC5 and additionally AC2, AC3 and AC6, however, AC2, AC5 and AC6 are not activated via Ca^2+^ and thus not involved in Ca^2+^-dependent cAMP signals as described in the present study (Brand et al., 2013; Brust et al., 2017; Onda et al., 2001). Soluble adenylyl cyclase AC10 is highly expressed in astrocytes in some brain regions such as the hippocampus and is Ca^2+^-activated (Choi et al., 2012; Jaiswal & Conti, 2003), hence it could mediate PE-induced cAMP rises. However, neither ST034307 nor NKY80 have been reported to inhibit AC10, and the complete inhibition of PE-evoked cAMP signals by combined ST034307 and NKY80 strongly suggests that not AC10, but AC1 and AC3 are the main Ca^2+^-activated adenylyl cyclases in olfactory bulb astrocytes. Immunohistochemical data supported the presence of AC1 and AC3 in OB astrocytes. While prior studies emphasized AC8 expression in the entire OB by in situ hybridization (Choi et al., 2012; Jaiswal & Conti, 2003; Muglia et al., 1999), our data suggests that AC1 and AC3 are more relevant in OB astrocytes and AC8 might be expressed in other cell types. While we observed AC8 immunostaining in the olfactory bulb, we did not find clear co-localization of AC8 and astrocyte markers, and lack of AC8-dependent cAMP signaling might be due to absence of AC8 in astrocytes.

## Conclusion and implications

Together, our findings provide a comprehensive view of NE-induced cAMP signaling in OB astrocytes and reveal receptor-specific and Ca^2+^-dependent as well as Ca^2+^-independent signaling pathways. NE activates astrocytic cAMP production via α_1_, α_2_, and β receptors, with distinct downstream mechanisms. Increases in cAMP evoked by β receptors were Ca^2+^-independent and employ the canonical G_S_ pathway, whereas PE-induced responses rely exclusively on intracellular Ca^2+^ signaling and require combined α_1A/D_ receptor activation for full activation of Ca^2+^-dependent AC1/AC3. Activation of α_2_ receptors also elicits both Ca^2+^ and cAMP rises, suggesting they employ the same signaling mechanism involving Ca^2+^-dependent stimulation of AC1 and AC3, however, when cAMP levels are elevated, α_2_ receptors inhibit cAMP production by G_i_. These results underscore the complex integration and tuning of cAMP and Ca^2+^ signaling in astrocytes and, in consideration of the importance of both Ca^2+^ and cAMP for metabolic interaction as well as synaptic plasticity, indicate that both second messenger systems strongly interact to optimize neuronal performance.

## Material and methods

### Animal handling

Mice were held in the animal facility of the Institute of Cell and Systems Biology (University of Hamburg, Hamburg, Germany) at a 12 h light/12 h dark cycle with food and water *ad libitum*. Animal handling and all experiments were approved by the local authorities (Behörde für Justiz und Verbraucherschutz, Lebensmittel und Veterinärwesen Hamburg; N010/2022) and followed German and European laws. To express the cAMP sensor Flamindo2 in astrocytes, C57Bl/6J wild type mice of both sexes at an age range from 4 to 16 weeks at the time of injection were used. They were injected with endotoxin-free recombinant adeno-associated viruses AAV^2/PhP.eB^hGFAP-Flamindo2 or AAV^2/PhP.AX^hGFAP-Flamindo2 (Odaka et al., 2014; Wendlandt et al., 2023). The capsid plasmid PhP-AX (Addgene #195218) was kindly provided by Vivianna Gradinaru and Xinhong Chen (Jang et al., 2023) and the capsid plasmid PhP.eB (Addgene #103005) was a gift from V. Gradinaru (Chan et al., 2017). All AAVs were produced at the vector facility of the University Medical Center Hamburg-Eppendorf (Hamburg, Germany). 70 µl virus suspension containing 1*10^11^ vg was injected i.v. into the retro-bulbar sinus under isoflurane-anesthesia (Fig. 1A). Experiments were performed 3 to 6 weeks after injection. For experiments with simultaneous cAMP and Ca^2+^ imaging, 1*10^11^ vg AAV^2/PhP.AX^ gfaABC1D-NES-jRCaMP1a (addgene # 171120, Lohr et al., 2021) was co-injected to image astrocytic Ca^2+^. For Ca^2+^ imaging without cAMP imaging, GLAST-CreERT2 x GCaMP6s^fl/wt^ mice (Chen et al., 2013; Mori et al., 2006) at an age range between 6 and 21 weeks were injected intraperitoneal three days in a row with tamoxifen (Carbolution Chemicals, St. Ingbert, Germany), 10 mg/ml in Miglyol (Caelo, Hilden, Germany) at 10 µl/g mouse weight. Imaging was conducted between 7 and 12 days after the first tamoxifen injection.

### Solutions and drugs

Artificial cerebrospinal fluid (ACSF) used during experiments contained (mM): 120 NaCl, 26 NaHCO_3_, 1 NaH_2_PO_4_, 2.5 KCl, 2.8 D-glucose, 2 CaCl_2_, 1 MgCl_2_. For the preparation of the OBs and brain slicing, Ca^2+^-reduced ACSF was used containing (in mM): 83 NaCl, 26.2 NaHCO_3_, 1 NaH_2_PO_4_, 2.5 KCl, 70 sucrose, 20 D-glucose, 0.5 CaCl_2_, 2.5 MgSO_4_. For experiments in Ca^2+^-free conditions (0Ca^2+^), ACSF was used containing (mM): 120 NaCl, 26 NaHCO_3_, 1 NaH_2_PO_4_, 2.5 KCl, 2.8 D-glucose, 3 MgCl_2_, 0.5 EGTA. All components were obtained from Carl Roth (Karlsruhe, Germany).

The following drugs were used: (R)-(−)-Phenylephrine hydrochloride (3-[(1R)-1-hydroxy-2-(methylamino)ethyl]phenol;hydrochloride; Abcam, Cambridge, United Kingdom), 2-APB (2-diphenylboranyloxyethanamine; Calbiochem, Darmstadt, Germany), BMY7378 (8-[2-[4-(2-methoxyphenyl)piperazin-1-yl]ethyl]-8-azaspiro[4.5]decane-7,9-dione;dihydrochloride; Hello Bio, Bristol, United Kingdom), CGP 55845 hydrochloride (benzyl-[(2S)-3-[[(1S)-1-(3,4-dichlorophenyl)ethyl]amino]-2-hydroxypropyl]phosphinic acid; hydrochloride; Hello Bio), Cyclopiazonic acid ((2R,3S,9R)-5-acetyl-4-hydroxy-8,8-dimethyl-7,16-diazapentacyclo[9.6.1.02,9.03,7.015,18]octadeca-1(17),4,11(18),12,14-pentaen-6-one; Hello Bio), D-AP5 ((2R)-2-amino-5-phosphonopentanoic acid; Alomone Labs, Jerusalem, Israel), Norepinephrine bitartrate (4-[(1*R*)-2-amino-1-hydroxyethyl]benzene-1,2-diol;2,3-dihydroxybutanedioic acid; Merck KGaA, Darmstadt, Germany), Forskolin ([(3R,4aR,5S,6S,6aS,10S,10aR,10bS)-3-ethenyl-6,10,10b-trihydroxy-3,4a,7,7,10a-pentamethyl-1-oxo-5,6,6a,8,9,10-hexahydro-2H-benzo[f]chromen-5-yl] acetate; Cayman Chemical, Ann Arbor, Michigan, USA), Gabazine (4-[6-imino-3-(4-methoxyphenyl)pyridazin-1-yl]butanoic acid; Abcam), HC-067047 (2-methyl-1-(3-morpholin-4-ylpropyl)-5-phenyl-N-[3-(trifluoromethyl)phenyl]pyrrole-3-carboxamide; Calbiochem), ICI 118,551 hydrochloride ((2R,3S)-1-[(7-methyl-2,3-dihydro-1H-inden-4-yl)oxy]-3-(propan-2-ylamino)butan-2-ol;hydrochloride; Tocris Bioscience, Bristol, United Kingdom), Isoprenaline hydrochloride (4-[1-hydroxy-2-(propan-2-ylamino)ethyl]benzene-1,2-diol;hydrochloride; Abcam), NBQX (disodium;6-nitro-7-sulfamoylbenzo[f]quinoxaline-2,3-diolate; Alomone Labs), NKY80 (2-amino-7-(furan-2-yl)-7,8-dihydro-6H-quinazolin-5-one; Tocris Bioscience), Prazosin hydrochloride ([4-(4-amino-6,7-dimethoxyquinazolin-2-yl)piperazin-1-yl]-(furan-2-yl)methanone;hydrochloride; Tocris Bioscience), Rauwolscine hydrochloride (methyl (1S,15S,18S,19S,20S)-18-hydroxy-1,3,11,12,14,15,16,17,18,19,20,21-dodecahydroyohimban-19-carboxylate;hydrochloride; Tocris Bioscience), RS100329 hydochloride (5-methyl-3-[3-[4-[2-(2,2,2-trifluoroethoxy)phenyl]piperazin-1-yl]propyl]-1H-pyrimidine-2,4-dione;hydrochloride;

Tocris Bioscience), ST 034307 (6-chloro-2-(trichloromethyl)chromen-4-one; Tocris Bioscience), tetrodotoxin citrate ((1R,5R,6R,7R,9S,11S,12S,13S,14S)-3-amino-14-(hydroxymethyl)-8,10-dioxa-2,4-diazatetracyclo[7.3.1.17,11.01,6]tetradec-3-ene-5,9,12,13,14-pentol; Hello Bio), Xylazine (N-(2,6-dimethylphenyl)-5,6-dihydro-4H-1,3-thiazin-2-amine;hydrochloride; Merck). Stock solutions of drugs were prepared as stated by the manufacturer. Stocks solutions were diluted in ACSF and applied via the perfusion system.

### Preparation of acute brain slices

Mice were anesthetized with isoflurane (5% v/v in oxygen) and decapitated. OBs were quickly dissected in chilled Ca^2+^-reduced ACSF. 220 µm sagittal slices of OBs (Fig. 1 B, C) were prepared using a vibratome (VT1200S, Leica, Bensheim, Germany), stored in ACSF at 30 °C for 30 min, and then kept at room temperature until the start of the experiment. Solutions were steadily gassed with carbogen (95% O_2_, 5% CO_2_).

### Confocal Imaging

After transferring slices to the recording chamber, imaging was performed with a confocal microscope (eC1, Nikon, Düsseldorf, Germany). Flamindo 2 was excited at 488 nm and emission detected from 500 to 530 nm. Images of astrocytes located in the glomerular and external plexiform layers (see Fig. 1C) (Lohr et al., 2014) were acquired at a rate of one frame every 5 s. Noradrenergic agonists were applied for 30 s via the perfusion system, using a peristaltic pump and a pump speed of 2.35 ml/min. Antagonists were incubated for 10 min before agonists were applied, except inhibitors of adenylyl cyclases (ACs) which were incubated for 30 min. Similar settings were used to image intracellular Ca^2+^ using GCaMP6s. For simultaneous imaging of cAMP and Ca^2+^, Flamindo2 was excited at 488 nm and jRCaMP1b was excited at 543 nm and emission collected between 553 and 618 nm. For H^+^ (pH) measurements, acute slices were incubated with 10 µM pHrodo Red AM and 100 µM PowerLoad (both from Invitrogen, Carlsbad, CA, USA) for 30 min and the same optical parameters were used for pHrodo Red imaging as for jRCaMP1b.

### Immunohistochemistry

Immunohistological staining was performed as described before (Beiersdorfer et al., 2020). Brains were fixed in 4 % Roti-Histofix (Roth, Karlsruhe, Germany) for 1 h and washed three times in PBS (in mM): 130 NaCl, 7 Na_2_HPO_4_, 3 NaH_2_PO_4_, pH adjusted to 7.4. OBs were sliced into 150 µm sagittal slices by a vibratome (VT1000, Leica, Bensheim, Germany). Antigen retrieval was performed in TE buffer (containing in mM: 10 Tris-HCl, 1 EDTA, 0.05 % Tween; pH adjusted to 9.0) at 95–98 °C for 15–20 min. Following a cooling period of 35–40 min, slices were incubated in blocking solution (5 % normal goat serum (NGS, Cell Signaling Technology) and 0.5 % Triton X-100 in PBS). After blocking, slices were incubated with primary antibodies for 48 h at room temperature diluted in primary antibody solution (blocking solution 1:10 in PBS), three times washed in PBS and subsequently incubated in secondary antibodies (dilution: 1:1000 in PBS) at 4 °C for 24 h. Afterwards slices were washed 3 X in PBS, and embedded on microscope slides (Thermo Fisher Scientific) with Shandon Immu-Mount (Thermo Fisher Scientific) and cover-slipped (Carl Roth). Images were captured by confocal microscope (eC1, Nikon, Düsseldorf, Germany) and processed using Fiji ImageJ (http://imagej.org) and GIMP2.0 (https://www.gimp.org). Schematic drawings were prepared with Biorender (www.biorender.com; license numbers GK28ZAAYYM, GP28ZAAVAO, ZS28ZAAMUV).

The following antibodies were used: Rabbit anti-ADCY1 (1:200; 55067-1-AP; Proteintech, Rosemont, Illinois, USA), rabbit anti-ADCY3 (1:200; 19492-1-AP; Proteintech), rabbit anti-ADCY8 (1:200; 55065-1-AP; Proteintech), rabbit anti-GFAP (1:500; Z0334 429-2; Dako, Agilent Technologies, Glostrup, Denmark), chicken anti-GFP (1:500; NB100-1614; Novus Biologicals, Centennial, Colorado, USA), rabbit anti-S100B (1:500; Z0311; Dako). Secondary antibodies: Goat anti-chicken Alexa 488 (1:1000; Invitrogen), goat anti-chicken Alexa 555 (1:1000; Invitrogen (part of Thermo Fisher Scientific, Carlsbad, California, USA), goat anti-rabbit Alexa 488 (1:1000; Invitrogen), goat anti-rabbit Alexa 555 (1:1000; Invitrogen). Nuclear stain: DAPI (5 µM; A1001; Applichem, Darmstadt, Germany).

### Data analysis and statistics

Imaging data was analyzed by marking cells as regions of interest (ROIs) in EZ-C1Viewer software (Nikon) and extracting data to Excel (Microsoft, USA) and Origin Pro 9.1 (OriginLab Corporation, Northampton, USA). The intensity of basal fluorescence F was set to 100 % and relative changes of fluorescence were measured (ΔF). Signals with relative changes below 3 % ΔF did not stand out significantly against noise and were excluded from further analysis. In contrast to GCaMP6s and pHrodo Red, which increase their fluorescence with an increasing concentration of the respective ion, an increase in the intracellular cAMP concentration is reflected by a decrease in Flamindo2 fluorescence (Odaka et al., 2014) and we depict fluorescence traces inverted (−ΔF) to better illustrate actual cAMP changes. Traces represent one single exemplary cell. All values are stated as mean values ± standard error of the mean. The number n corresponds to the number of analyzed cells. Each set of experiments was carried out on brain slices prepared from at least three animals. A sigmoidal fit was used to analyze the dose-response relationship of NE-evoked cAMP transients (Origin Pro 9.1). The following tests were used for evaluation of statistical significance: Outliers were identified and dismissed by Grubbs test. Wilcoxon signed rank test was used for paired data, Mann-Whitney-U test for independent data. For comparisons involving more than two groups, the Kruskal-Wallis ANOVA followed by Dunńs post hoc test was applied for independent data, and the Friedmann ANOVA followed by Wilcoxon signed-rank post hoc test for paired data. Significant differences were assumed with error probabilities p < 0.05 (*p < 0.05, **p < 0.01, ***p < 0.001).

## Acknowledgements

We thank AC Rakete (University of Hamburg) for excellent technical assistance. Financial support by the Deutsche Forschungsgemeinschaft is greatfully acknowledged (SFB 1328 TP A07, 404539526, and A16, 404644007).

## Contributions

Study design: JS, DH, CEG, CL. Methodology: JS, OC, DH, CEG, CL. Experiments: JS, AB, FLS, MN. Data analysis: JS, AB, FLS, MN. Writing and figure design: JS, AB, CEG, CL. All authors edited and approved the manuscript.

## Supplementary Materials

**Fig. s1.**
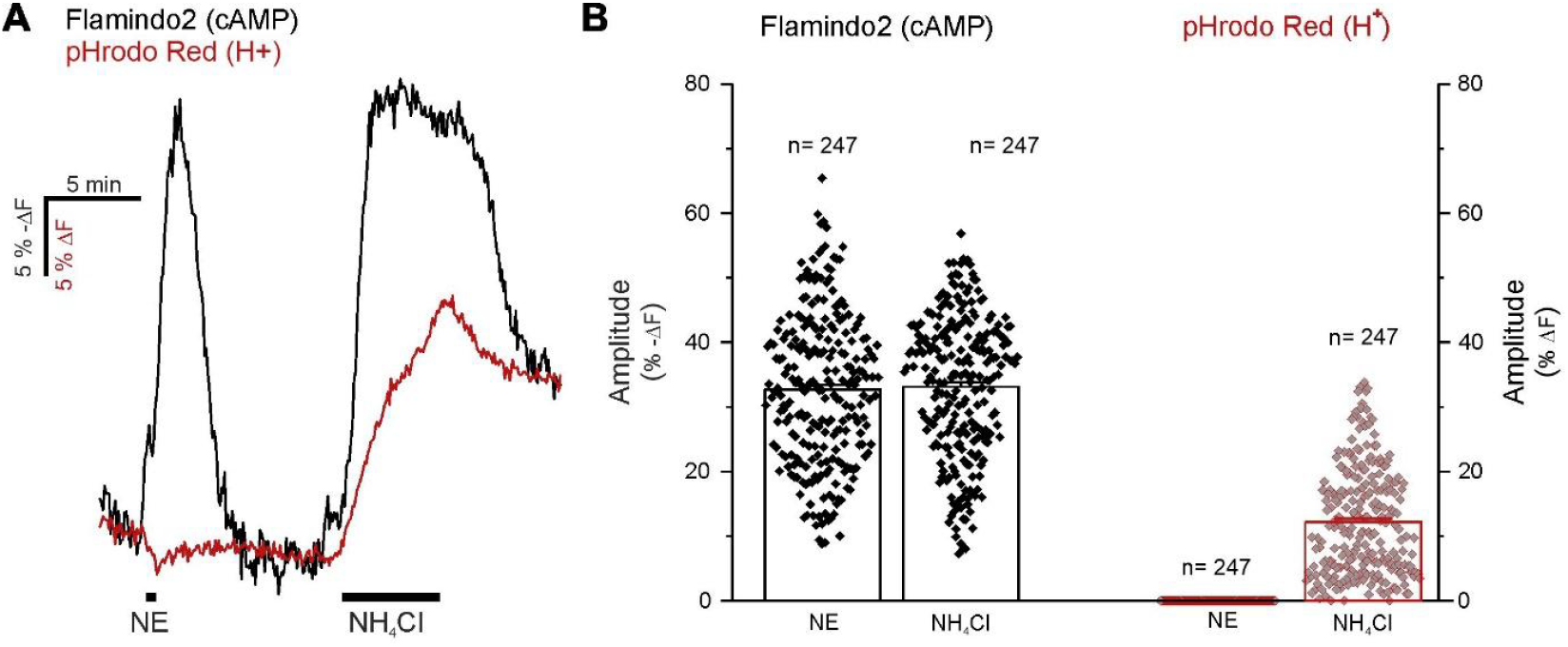
Effect of pH changes on Flamindo2 fluorescence. (**A**) Changes in cAMP (black) and H^+^ (red) concentrations induced by norepinephrine (NE) and NH_4_Cl. Note that the Flamindo2 trace is inverted (−ΔF) and Flamindo2 fluorescence decreases upon acidification by NH_4_Cl. (**B**) Quantification of cAMP and H^+^ changes. The results demonstrate pH sensitivity of Flamindo2, however, application of NE fails to evoke pH shifts, indicating that NE-evoked changes in Flamindo2 fluorescence are not pH-dependent. Data from 4 mice.

**Fig. s2.**
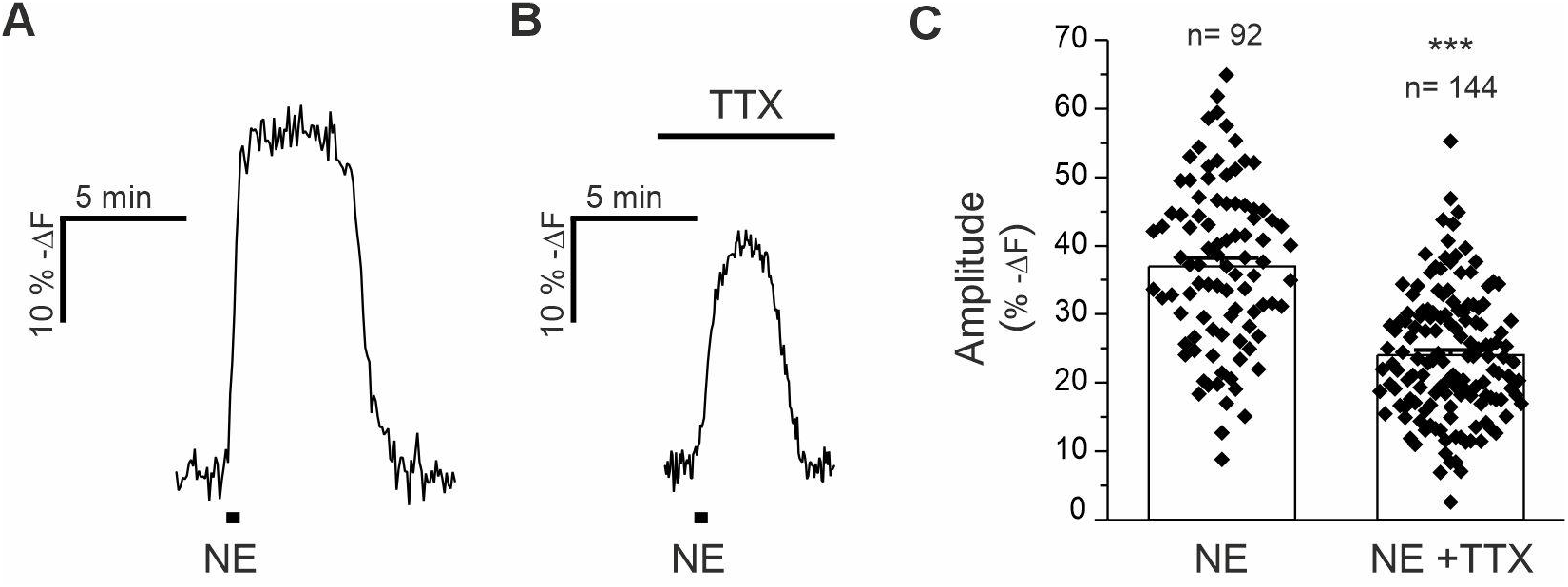
Tetrodotoxin (TTX) reduces NE-evoked cAMP responses in OB astrocytes. (A) Bath application of norepinephrine (NE, 10 µM) for 30 s resulted in transient increases in cAMP in the absence (control) and in (B) the presence of 0.5 µM TTX. (C) NE-evoked increases in cAMP were significantly reduced by TTX. ***p < 0.001, Mann-Whitney U-test; data from 6 mice for control (NE), 6 mice for TTX (NE + TTX).

**Fig. s3.**
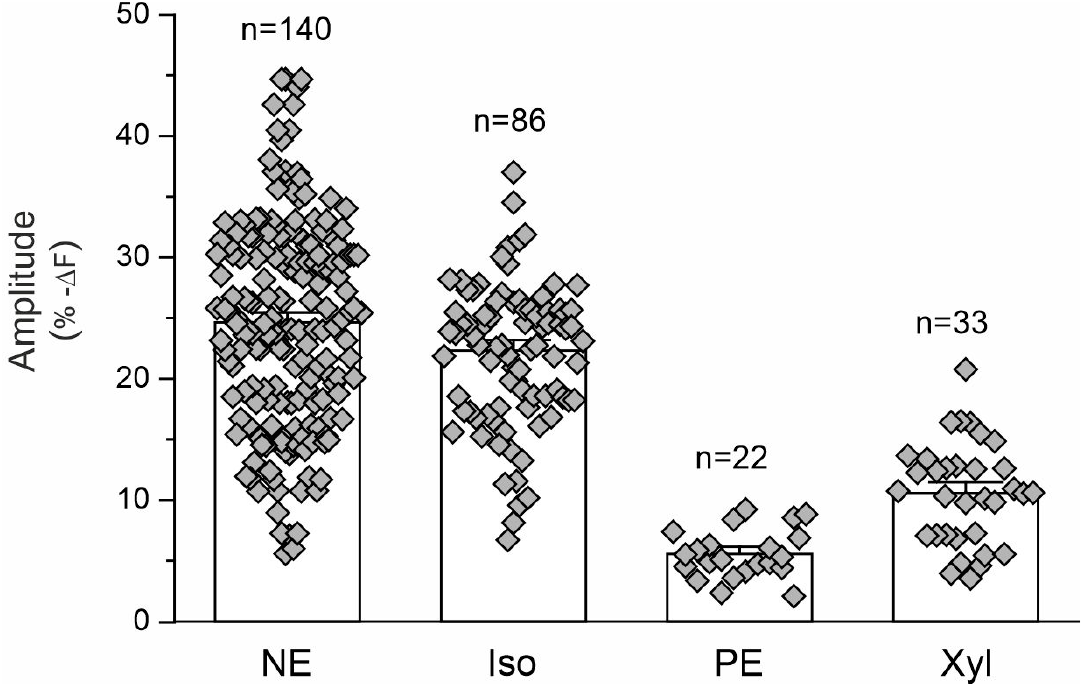
Amplitudes of cAMP responses evoked by various adrenergic agonists in OB astrocytes. Bath application of norepinephrine (NE, 10 µM; data from 6 mice), isoprenaline (Iso, 100 µM; data from 3 mice), phenylephrine (PE, 100 µM; data from 4 mice) and xylazine (Xyl, 80 µM; data from 3 mice) evoked increases in the cAMP concentration.

**Fig. s4.**
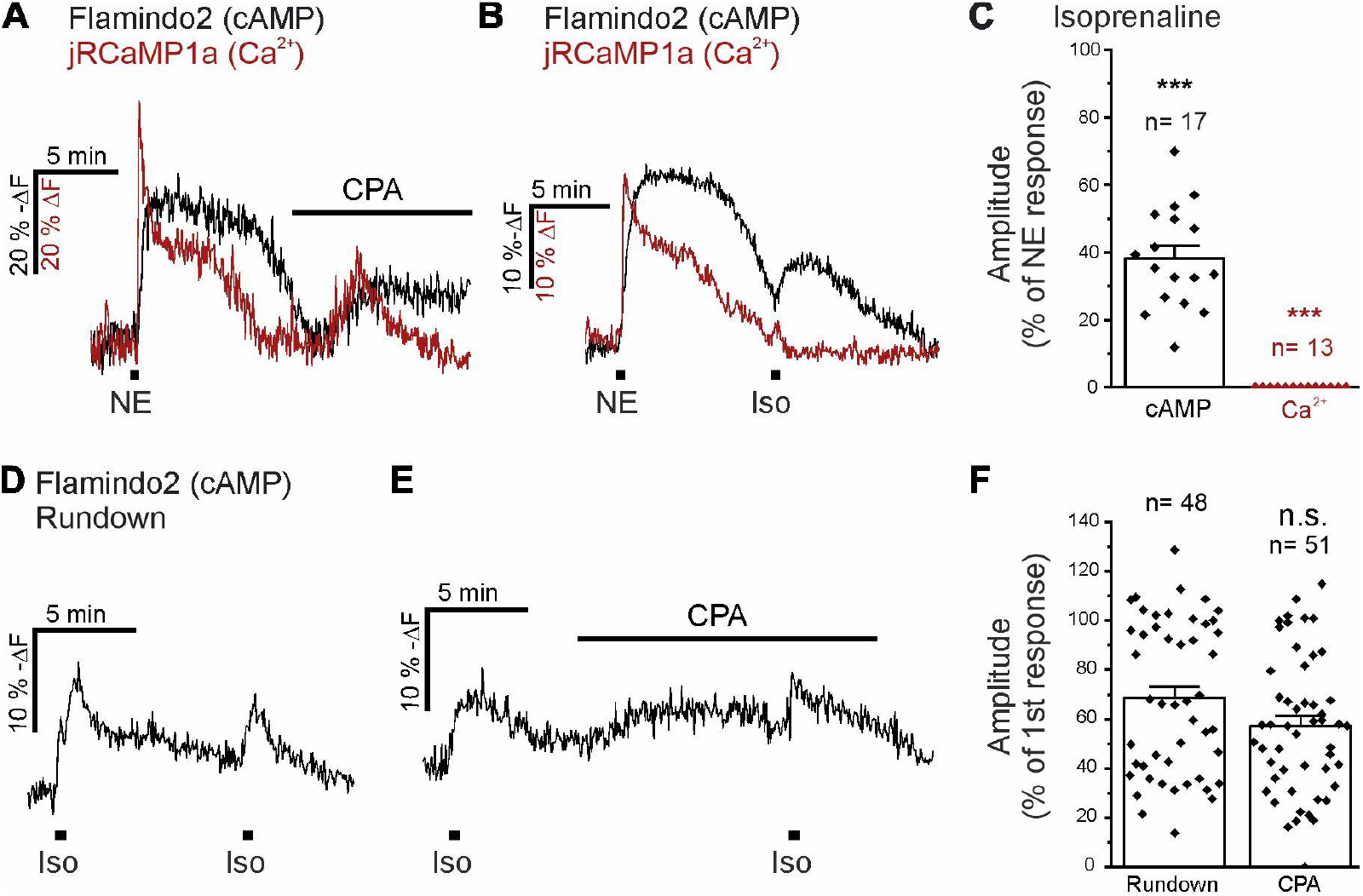
Cyclopiazonic acid and isoprenaline induced cAMP signals in olfactory bulb astrocytes. (**A**) Depletion of intracellular Ca^2+^ stores by cyclopiazonic acid is visible by a moderate increase in jRCaMP1a fluorescence (red trace) and accompanied by an increase in Flamindo2 fluorescence (black trace), indicating a rise in cAMP. Norepinephrine (NE, 10 µM) was applied as a control to test for vital Ca^2+^ and cAMP signaling. (**B**) NE evoked both cAMP (black trace) and Ca^2+^ signals (red trace), whereas isoprenaline (Iso, 100 µM) induced only cAMP signals. (**C**) Analysis of the Iso-induced cAMP and Ca^2+^ signals, normalized to the amplitude of the NE-evoked response which was set to 100 %. ***p < 0.001, Mann-Whitney-U test. Rundown control: Data from 3 mice Flamindo2, 3 mice Flamindo2 + jRCaMP1a; Iso: data from 2 mice Flamindo2 + jRCaMP1a. (**D**) cAMP signals evoked by repetitive application of Iso as a rundown experiment. (**E**) Effect of 20 µM cyclopiazonic acid (CPA) on Iso-evoked cAMP signals. (**F**) In the presence of CPA, Iso-evoked cAMP responses were not significantly altered compared to the corresponding rundown experiment. n.s. not significant, Mann-Whitney-U test; rundown: data from 3 mice; CPA: data from 3 mice.

**Fig. s4a.**
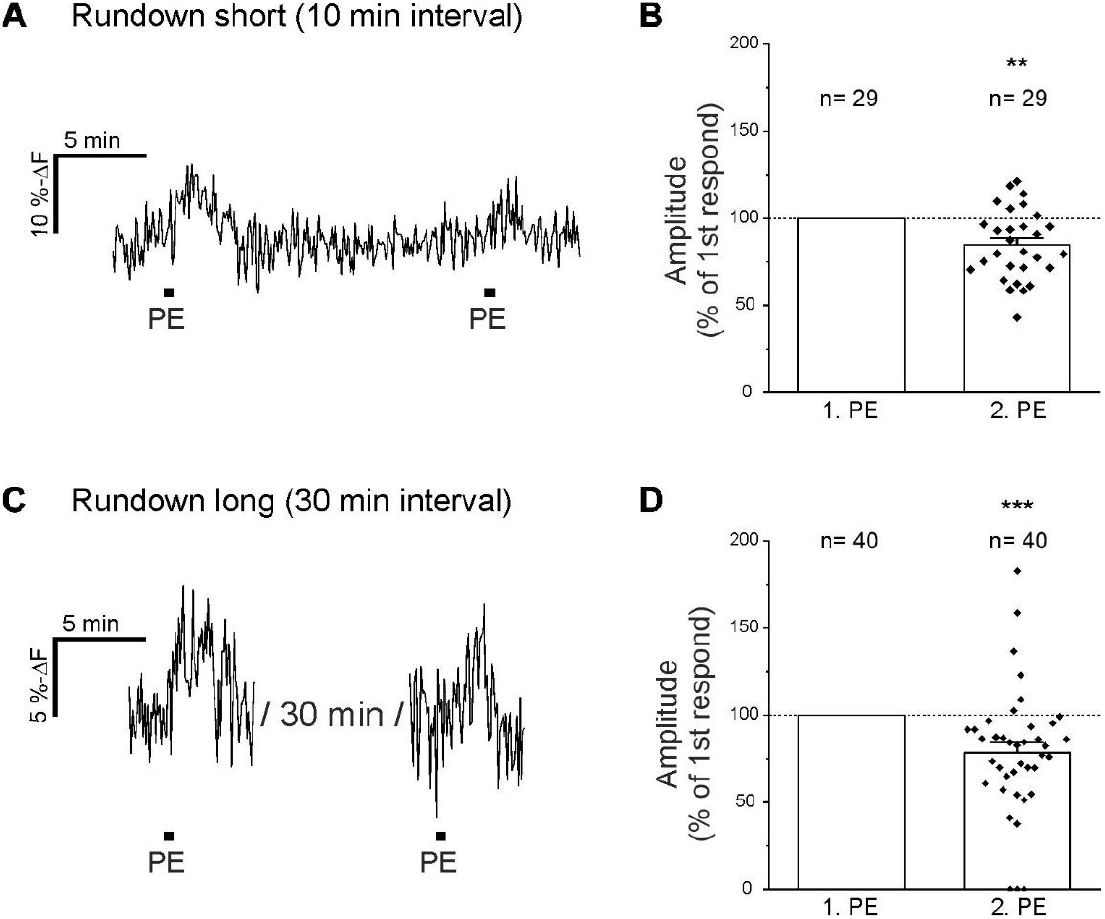
Phenylephrine rundown experiments. (**A**) Phenylephrine (PE)-induced cAMP signals evoked by two PE applications with intervals of 10 minutes (rundown short). **(B)** A second application of PE evoked significantly smaller responses compared to the first application. **p < 0.01, Mann-Whitney-U test, data from 3 mice. (**C**) PE applications with intervals of 30 minutes (rundown long). **(B)** A second application of PE evoked significantly smaller responses compared to the first application. ***p < 0.001, Mann-Whitney-U test; data from 7 mice.

**Fig. s5.**
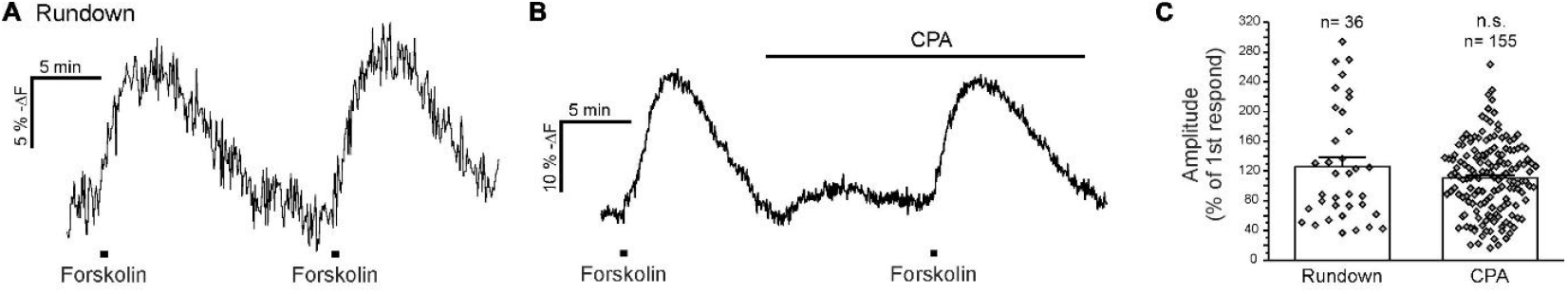
Cyclopiazonic acid has no effect on adenylyl cyclase activity. (**A**) Astrocytic cAMP signals evoked by repetitive application of 3 µM forskolin. (**B**) Forskolin-evoked cAMP transients before and in the presence of 20 µM cyclopiazonic acid (CPA). (**C**) CPA has no significant effect on forskolin-evoked cAMP increases n.s. not significant, Mann-Whitney-U test, data from 4 mice for rundown, 5 mice for CPA.

**Fig. s6.**
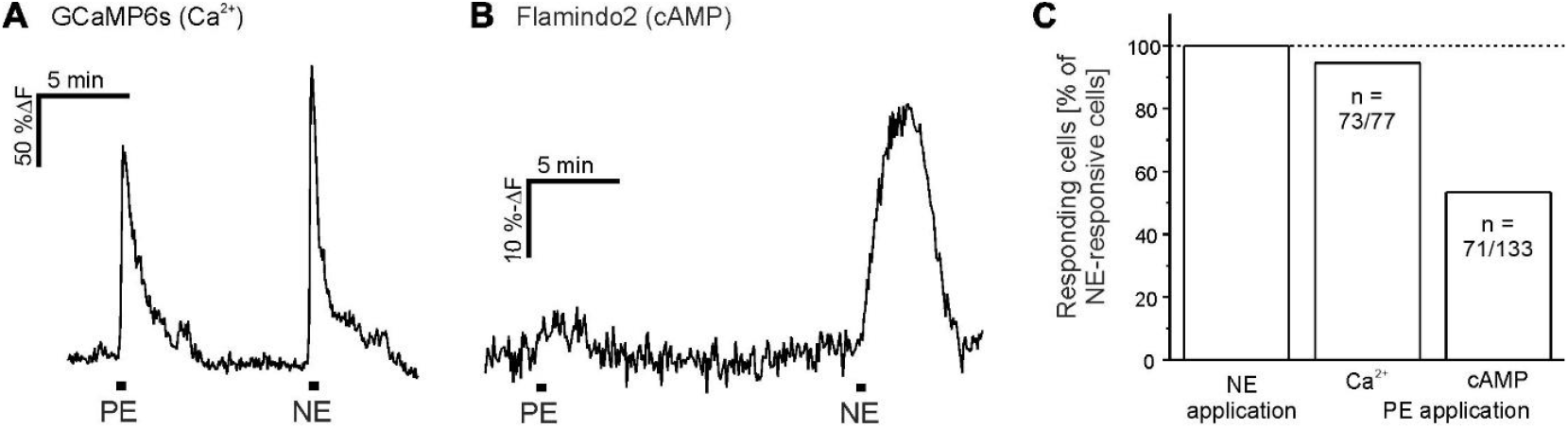
Ca^2+^ and cAMP signaling induced by phenylephrine and norepinephrine. (**A**) Phenylephrine (PE)- and norepinephrine (NE)-induced Ca^2+^ and (**B**) cAMP signals. (**C**) Fraction of responding astrocytes. While virtually all astrocytes responded to NE and PE with Ca^2+^ transients, only 53 % of astrocytes (71 out of 133) responded to PE application with an increase in cAMP. Data from 3 mice for Ca^2+^, 3 mice for cAMP.

**Fig. s7.**
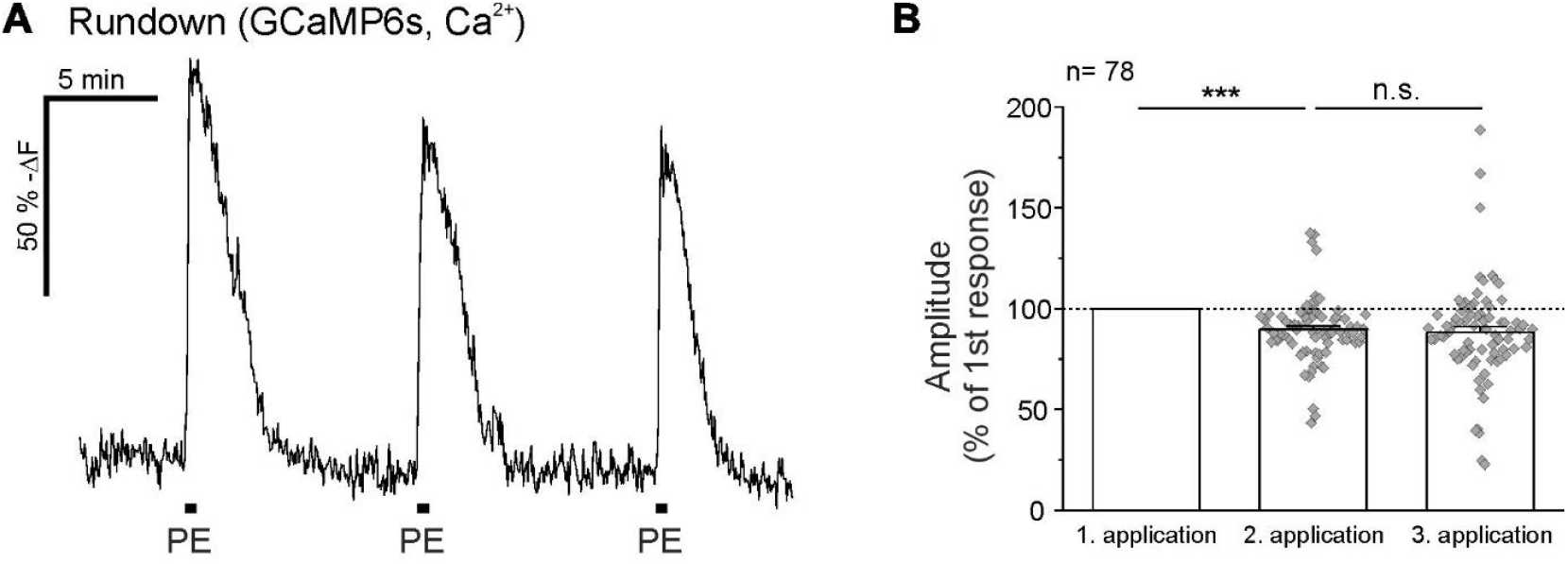
Rundown of three phenylephrine (PE)-induced Ca^2+^ signals. **(A)** Ca^2+^ signals in olfactory bulb astrocytes evoked by repetitive PE-applications. **(B)** Analysis of the Ca^2+^ signal amplitudes, normalized to the amplitude of the first response. n.s.=not significant, ***, p < 0.001. Friedmann ANOVA and Wilcoxon signed-rank post hoc test, data from 4 mice.

**Table s1.** Amplitudes of norepinephrine-induced cAMP signals in astrocytes at concentrations ranging from 1–30 µM (raw values for dose-response curve in Fig.1J). Data from 3 mice.

| Concentration norepinephrine | Mean amplitudes ± SEM (% $-\Delta\text{F}$) | n |
| --- | --- | --- |
| 1 $\mu\text{M}$ | 6.1 ± 0.8 | 88 |
| 3 $\mu\text{M}$ | 23.7 ± 1.1 | 88 |
| 10 $\mu\text{M}$ | 27.4 ± 0.9 | 88 |
| 30 $\mu\text{M}$ | 31.0 ± 0.9 | 88 |

**Table s2.** Amplitudes of phenylephrine-induced cAMP signals in astrocytes at concentrations ranging from 1–300 µM (raw values for dose-response curve in Fig. 4B). Data from 6 mice.

| **Concentration phenylephrine** | **Mean amplitudes ± SEM (% $-\Delta F$)** | **n** |
| --- | --- | --- |
| 1 μM | 3.7 ± 0.5 | 61 |
| 3 μM | 6.3 ± 0.3 | 84 |
| 10 μM | 6.6 ± 0.2 | 112 |
| 30 μM | 7.7 ± 0.4 | 80 |
| 100 μM | 8.3 ± 0.5 | 57 |
| 300 μM | 8.7 ± 0.7 | 29 |

## Notes

### Competing Interest Statement

The authors have declared no competing interest.

